# Establishing synthetic ribbon-type active zones in a heterologous expression system

**DOI:** 10.1101/2024.03.18.585517

**Authors:** Rohan Kapoor, Thanh Thao Do, Niko Schwenzer, Arsen Petrovic, Thomas Dresbach, Stephan E. Lehnart, Rubén Fernández-Busnadiego, Tobias Moser

## Abstract

Encoding of several sensory modalities into neural signals is mediated by ribbon synapses. The synaptic ribbon tethers synaptic vesicles at the presynaptic active zone (AZ) and may act as a super-scaffold organizing AZ topography. Here we employed a synthetic biology approach to reconstitute structures mimicking ribbon-type AZs in HEK293 cells for probing minimal molecular requirements and studying presynaptic Ca^2+^ channel clustering. Co-expressing a membrane-targeted version of the AZ-protein Bassoon and the ribbon core protein RIBEYE, we observed structures recapitulating basic aspects of ribbon-type AZs, which we call *synthetic* ribbons or *SyRibbons*. Super-resolution STED microscopy and cryo-correlative electron tomography revealed *SyRibbons* were similar to native ribbons at AZs of cochlear inner hair cells in shape and size. *SyRibbons* with Ca^2+^ channel clusters formed upon additional expression of Ca_V_1.3 Ca^2+^ channels and RIM-binding protein 2 (RBP2). Ca_V_1.3 Ca^2+^ channel clusters associated with *SyRibbons* were larger than ribbonless Ca_V_1.3 Ca^2+^ channels clusters and functional analysis by Ca^2+^-imaging in combination with patch clamp showed partial confinement of the Ca^2+^ signal at *SyRibbons*. In summary, we identify Ca^2+^ channels, RBP, membrane-anchored Bassoon, and RIBEYE as minimal components for reconstituting a basic ribbon-type AZ. *SyRibbons* might complement animal studies on molecular interactions of AZ proteins.

**Significance Statement:** Encoding of sensory information in our eyes and ears builds on specialized ribbon synapses of sensory cells. Elucidating the molecular underpinning of their fascinating structure and function is an ongoing effort to which we add a bottom-up reconstitution approach in cultured cells. Aiming to recapitulate basic properties of ribbon-type presynaptic active zones of cochlear inner hair cells, we identified a minimal set of proteins that assemble in cellular nanodomains, structurally and functionally alike active zones. While not yet reconstituting synaptic vesicle exocytosis, we consider the established synthetic ribbon-type active zones a valuable platform for studying molecular interactions of active zone proteins. We expect the approach to complement and reduce experiments on native ribbon synapses asserted from animals.

## Introduction

Synapses of sensory receptor cells of the inner ear and the retina are hallmarked by the presence of an electron-dense structure called the synaptic ribbon, which tethers synaptic vesicles (SVs) at the presynaptic active zone (AZ, [Moser *et al*, 2019; Matthews & Fuchs, 2010]). Proposed functional roles of the ribbon include (i) its role as a SV replenishment machine for tireless neurotransmission (Bunt, 1971; Holt *et al*, 2004; Khimich *et al*, 2005; Frank *et al*, 2010; Snellman *et al*, 2011; Graydon *et al*, 2011; Vaithianathan *et al*, 2016), (ii) “super-scaffold” regulating the abundance, topography, and function of AZ players such Ca^2+^ channels and release sites and their tight coupling (Khimich *et al*, 2005; Frank *et al*, 2010; Wong *et al*, 2014; Maxeiner *et al*, 2016; Jean *et al*, 2018; Grabner & Moser, 2021), and iii) coordination of SV release (Heidelberger *et al*, 1994; Glowatzki & Fuchs, 2002; Singer *et al*, 2004; Edmonds, 2004; Mehta *et al*, 2013). Yet, our molecular understanding of how these postulated functions are executed by synaptic ribbons is still limited.

Likewise, the clustering of Ca^2+^ channels at the AZ, how it is varied among different or even the same synapse types and how the ribbon contributes towards this, remain active areas of research (Wichmann & Kuner, 2022; Moser *et al*, 2023). For example, ribbon synapses of mature inner hair cells (IHC) assemble Ca_V_1.3 voltage-gated Ca^2+^ channels at the base of the ribbon, but vary in shape, size and channel complement of the Ca_V_1.3 clusters (Frank *et al*, 2010; Wong *et al*, 2014; Neef *et al*, 2018) partially dependent on the position within the IHC (Ohn *et al*, 2016; Özçete & Moser, 2021). IHC synapses lacking AZ-anchored ribbons in Bassoon mutant mice showed fewer Ca^2+^ channels with altered topography (Frank *et al*, 2010; Neef *et al*, 2018). While genetic disruption of Ca^2+^ channel-tethering AZ proteins RIM2 (Jung *et al*, 2015) and RBP2 (Krinner *et al*, 2017) reduced the abundance of Ca^2+^ channels at IHC synapses, their topography retained the typical stripe-like shape of Ca^2+^ channel clusters. Constitutive deletion of RIBEYE, the core scaffold protein of the synaptic ribbon, provided so far the most direct test of the scaffolding role of the ribbon (Maxeiner *et al*, 2016; Jean *et al*, 2018; Becker *et al*, 2018; Grabner & Moser, 2021). Here, ribbon-less synapses of IHCs showed an altered Ca^2+^ channel topography with a more spatially widespread presynaptic Ca^2+^ signal but intact Ca^2+^ channel complement (Jean *et al*, 2018). Moreover, the voltage-dependence of activation as well as the inactivation of Ca^2+^ channels appeared to be altered, suggesting a possible role of the synaptic ribbon in regulating Ca^2+^ channel physiology (Jean *et al*, 2018). Changes in Ca^2+^ channel physiology upon deletion of RIBEYE were also found for Ca_V_1.4 at ribbon-less rod photoreceptor AZs (Grabner & Moser, 2021). Finally, IHCs lacking piccolino, a ribbon-synapse specific isoform of the AZ protein piccolo, show AZs with smaller synaptic ribbons and altered clustering, yet normal complement of Ca^2+^ channels (Michanski *et al*, 2023).

So how does the synaptic ribbon tune into the orchestra of AZ proteins that collectively organize Ca^2+^ channel abundance, topography, and function? To what extent can the effects observed upon disruption of Bassoon be attributed to the concomitant loss of the ribbon? These questions seem even more relevant given that, so far to our knowledge, no direct interactions between Bassoon (Frank *et al*, 2010) or RIBEYE and Ca_V_1.3 Ca^2+^ channel have been described. Here, we adopted a bottom-up synthetic biology approach to assemble a minimal presynaptic protein machinery required to bring together RIBEYE and Ca^2+^ channels in a ‘synapse-naïve’ expression system to assess the role of the synaptic ribbon in regulating Ca^2+^ channel clustering and physiology. Similar approaches have previously been employed for reconstituting aspects of conventional synapses, for instance in “hemisynapses” between co-cultured neurons and non-neuronal cells overexpressing presynaptic or postsynaptic components (for review see Craig *et al*, 2006). More recently, Munc13-1 supra-molecular assemblies which recruit release machinery proteins were reconstituted (Sakamoto *et al*, 2018).

Expression of RIBEYE in synapse-naïve cell lines such as COS-7 cells led to large cytosolic assemblies (Schmitz *et al*, 2000). Ribbon-like electron dense structures with vesicles have also been reported upon heterologous RIBEYE expression in retinal progenitor cell-line (R28, [Magupalli *et al*, 2008]). However, these cells might not be considered synapse-naïve and the presence and function of Ca^2+^ channels at synthetic ribbon synapses remained to be studied. The need for co-expressing at least three Ca^2+^ channel subunits (Ca_V_α_x_, Ca_V_β_x_ and Ca_V_α_2_8_x_) in synapse-naïve cell lines renders acute co-expression of AZ multidomain proteins cumbersome. Here we took advantage of synapse-naïve Human Embryonic Kidney 293 (HEK293) cells stably expressing inducible Ca_V_1.3α_1_, which is the predominant subtype of voltage-gated Ca^2+^ channels at IHC ribbon synapses (Brandt *et al*, 2003; Platzer *et al*, 2000; Dou *et al*, 2004), as well as constitutively expressed Ca_V_β_3_ and Ca_V_α_2_8_1_. We co-expressed RIBEYE along with membrane targeted Bassoon and observed structures with striking resemblance to IHC synaptic ribbon-type AZs nicknamed ‘*SyRibbons*’. We characterized the structure and function of these synthetic ribbon-type AZs and identified Bassoon, RBP2, RIBEYE and Ca_V_1.3 channels as the minimal components required for assembling a basic ribbon-type AZ. We observed larger Ca_V_1.3 channel clusters near *SyRibbons* and with partially localized Ca^2+^ influx upon stimulation. Our results support the role of synaptic ribbons in promoting the formation of Ca_V_1.3 channel micro-clustering. Synthetic ribbon-type AZs offer a novel approach for functionally studying protein-protein interaction and will likely complement, refine and reduce experiments on native ribbon synapses.

## Results

### Membrane-targeted Bassoon recruits RIBEYE to the cell membrane in HEK293 cells where structures mimicking inner hair cell synaptic ribbons are formed

HEK293 cells have the advantage of not expressing the synaptic machinery components studied here. This provides a clean background for reconstituting synthetic synapses from a minimal set of synaptic proteins. Our first step towards assembling a ribbon-type AZ in a heterologous expression system was to express RIBEYE, the core scaffold protein of the synaptic ribbon, and target it to the cell membrane. We performed transient transfection of a RIBEYE construct with a C-terminal EGFP tag in HEK293 cells and observed large spherical clusters of RIBEYE that appeared largely cytosolic (Fig. 1Aii), in contrast to a diffuse cytosolic distribution when merely expressing EGFP (Fig. 1Ai). These RIBEYE clusters form due to self-assembling properties of RIBEYE via multiple sites of homophilic interaction as have been demonstrated before in several cell lines. They do not colocalize with the endoplasmic reticulum (ER), Golgi apparatus, or lysosomes and hence, are unlikely to reflect RIBEYE entrapped in protein trafficking pathways or to represent degradation products of overexpressed RIBEYE (Supplementary Fig. 1A, B and C).

**Figure 1.**
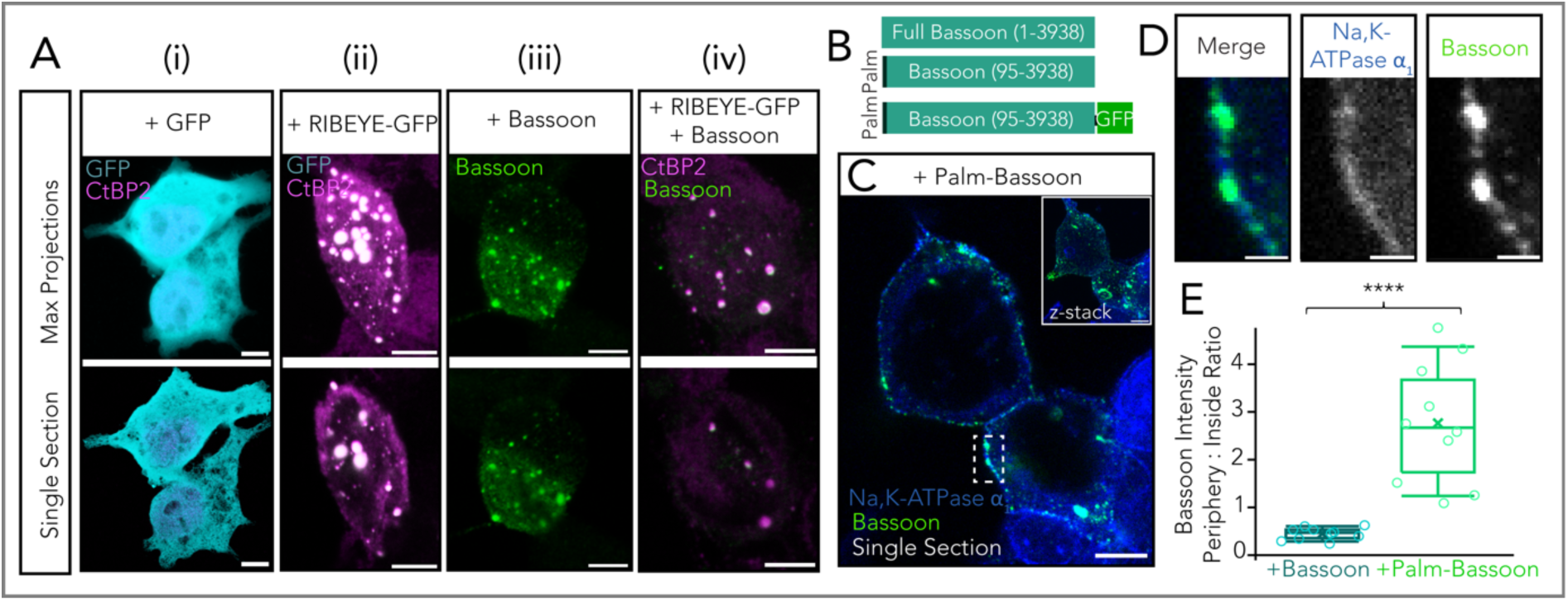
Membrane targeting of Bassoon using a palmitoylation consensus sequence. **A** Representative confocal images of HEK293 cells transfected with (i) GFP only (CtBP2/RIBEYE in magenta labeling nuclei, GFP in cyan), (ii) RIBEYE-GFP (CtBP2/RIBEYE in magenta, GFP in cyan), (iii) Bassoon (green) and (iv) RIBEYE-GFP and Bassoon. Note that antibodies against RIBEYE B-domain and the nuclear transcription factor CtBP2 result in similar staining patterns as CtBP2 is transcribed from the same gene as RIBEYE and is identical to the RIBEYE-B domain except for the first 20 N-terminal amino acids. Upper panel shows maximum projections and lower panel shows exemplary single sections from confocal stacks. Scale bar = 5 µm. **B** Schematic of construct for membrane targeting of Bassoon. The first 95 amino acids from full-length Bassoon were replaced with a palmitoylation consensus sequence from GAP43. Constructs without and with a C-terminal GFP tag were used, as depicted. **C** Sample confocal image (single section) showing membrane targeted palm-Bassoon (green) expressed in HEK293 cells appears as puncta distributed along the periphery of the cell, marked by Na, K-ATPase α_1_(blue). Inset shows maximal projection of confocal section. Scale bar = 5µm. **D** Zoom-in from **(C)** shows colocalisation of palm-Bassoon with membrane marker Na, K-ATPase α1. Scale bar = 1µm. **E** Quantification of Bassoon signal intensity at periphery *vs* inside of cell. Cells expressing palm-Bassoon (N = 10 cells) clearly show a higher peripheral distribution compared to cells expressing full-length Bassoon (N = 10 cells), *\*\*\*\*P* < 0.0001, Mann-Whitney-Wilcoxon test. Overlaid data points represent individual cells, crosses represent the mean values, central band indicates the median, whiskers represent 90/10 percentiles, and boxes represent 75/25 percentiles.

Next, for membrane targeting of these cytosolic RIBEYE clusters, we co-expressed the multidomain cytomatrix of the active zone protein Bassoon (Fig. 1Aiii). Prior work on the molecular underpinnings of ribbon synapses had identified Bassoon (tom Dieck *et al*, 1998) to critically contribute to anchoring the synaptic ribbon to the AZ membrane (Khimich *et al*, 2005; Dick *et al*, 2003; tom Dieck *et al*, 2005). Co-expression of full-length Bassoon along with RIBEYE in HEK293 cells showed colocalizing clusters of the two proteins that, however, remained largely cytosolic (Fig. 1Aiv).

Next, for plasma membrane-targeting of Bassoon, we generated a construct by removing the first 95 N-terminal amino acids of Bassoon and replacing these with a palmitoylation consensus sequence of the neuronal Growth Associated Protein 43 (GAP43). We refer to this as ‘palm-Bassoon’ throughout, and we used constructs with and without a C-terminal EGFP tag (Fig. 1B). Expression of either of these palm-Bassoon constructs in HEK293 cells showed comparable immunofluorescence patterns with Bassoon puncta spread across the periphery of the cell, largely colocalizing with the endogenously expressed plasma membrane-standing Na, K-ATPase α_1_ (data representative of 3 transfections, Fig. 1C, D). Palm-Bassoon does not appear to localize in the ER, Golgi apparatus or lysosomes (Supplementary Fig. 1D, E and F). Comparing the ratio of Bassoon signal intensity at the periphery *versus* inside of the cell in randomly selected single sections from confocal stacks of palm-Bassoon and Bassoon transfected cells (N = 10 cells, 3 transfections per group, Fig. 1E) demonstrated a higher Bassoon signal intensity at the periphery of cells expressing the palm-Bassoon construct (*\*\*\*\*P* < 0.0001, Mann-Whitney-Wilcoxon test), implying successful plasma membrane targeting of Bassoon.

Next, we co-expressed palm-Bassoon and RIBEYE in HEK293 cells and observed colocalizing RIBEYE and Bassoon immunofluorescent puncta at the periphery of cells (7 transfections; Fig. 2Ai). Closer inspection of these immunofluorescent puncta with STimulated Emission Depletion (STED) nanoscopy (Fig. 2Aii) revealed discrete structures typically consisting of ellipsoidal RIBEYE clusters juxtaposing on top of plate-like palm-Bassoon structures which seemingly anchor the RIBEYE clusters to the plasma membrane. We found the morphology of the RIBEYE + palm-Bassoon structures to be strikingly reminiscent of the arrangement of the two proteins in IHC ribbon synapses, where an ellipsoid/spherical synaptic ribbon composed of RIBEYE is found seated on a Bassoon plate that anchors it to the presynaptic AZ membrane (e.g. Wong *et al*, 2014; Michanski *et al*, 2019, 2023) (Fig. 2B, i and ii; data from Michanski *et al*, 2023).

**Figure 2.**
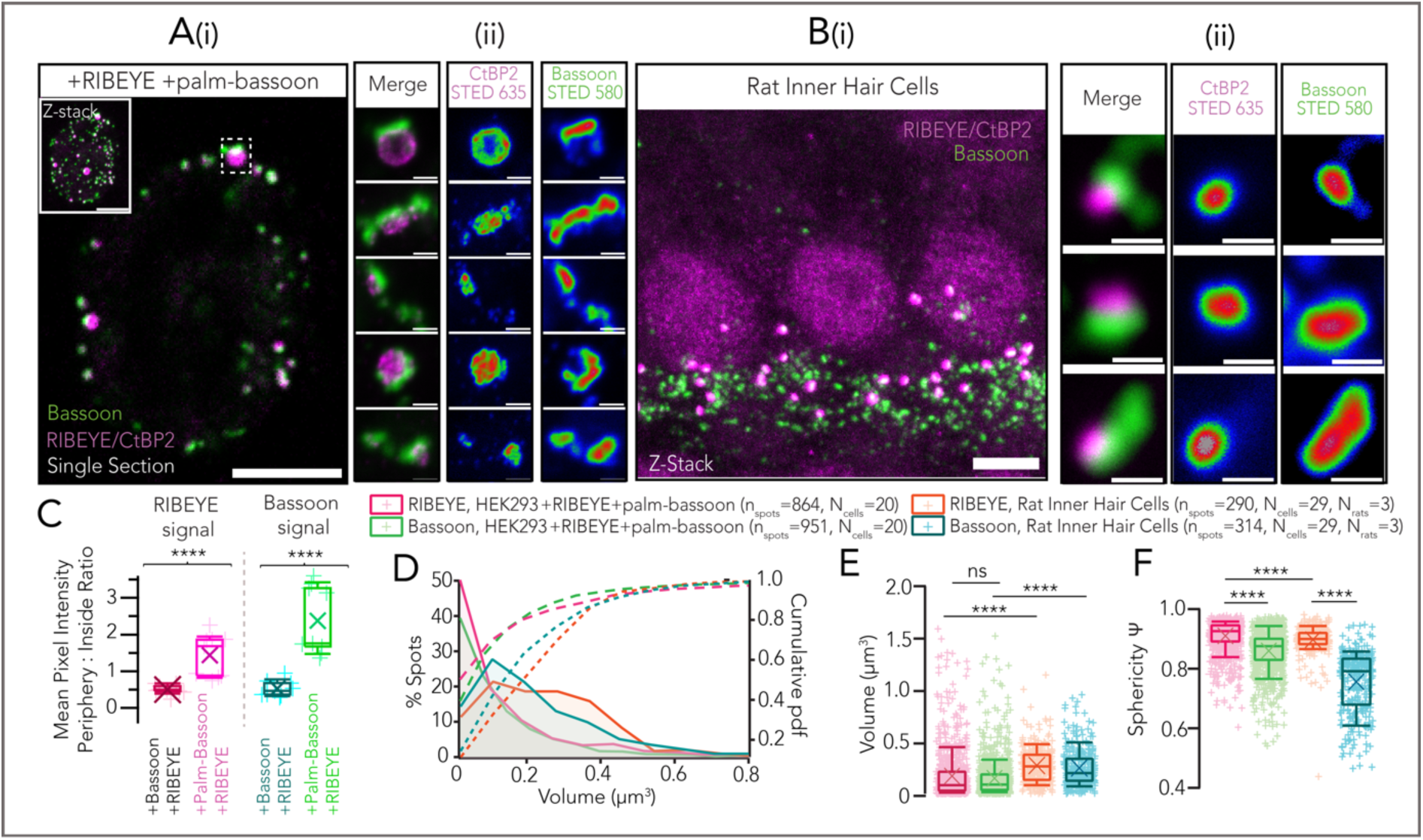
Co-expression of RIBEYE with palm-Bassoon results in ribbon-type AZ-like structures. A (i) Representative confocal image (single section) of a HEK293 cell transfected with RIBEYE-GFP (magenta) and palm-Bassoon (green). Co-expression of RIBEYE with palm-Bassoon targets RIBEYE to the cell membrane. Inset shows maximum projection. Scale bar = 5µm. (ii) Exemplary 2-D STED images for RIBEYE – palm-Bassoon juxtapositions acquired from cells as shown in (i); Scale bar = 500nm; individual channels have been depicted with an intensity-coded look-up table with warmer colors indicating higher intensity. B **(i)** Representative maximum projection of confocal sections of apical organ of Corti from a Wistar rat (postnatal day 18); data as published in Michanski *et al*, 2023, stainings were for CtBP2/RIBEYE (magenta; labeling synaptic ribbons and IHC nuclei) and Bassoon (green; spots juxtaposing with ribbons represent IHC AZs, spots not juxtaposing with ribbons represent efferent synapses formed by lateral olivocochlear neurons onto SGN boutons). Scale bar = 5µm. **(ii)** Juxtaposing RIBEYE and Bassoon spots imaged in 2-D STED and confocal mode respectively. Scale bar = 500nm. Note the striking resemblance to reconstituted RIBEYE + palm-Bassoon structures in HEK293 cells as shown in **(Aii)**. STED images are from 3 sample transfections, representative of 7 total transfections; individual channels have been depicted with an intensity-coded look-up table with warmer colors indicating higher intensity. C Quantification of RIBEYE and Bassoon signal intensity at periphery *vs* inside of cell shows a higher peripheral distribution of RIBEYE and Bassoon in HEK293 + RIBEYE + palm-Bassoon cells (N=9 cells) as compared to HEK293 + RIBEYE + Bassoon cells (N=9 cells); *\*\*\*\*P* < 0.0001, Mann-Whitney-Wilcoxon test. Overlaid data points represent individual cells, crosses represent mean values, central band indicates the median, whiskers represent 90/10 percentiles, and boxes represent 75/25 percentiles. D Distribution of volumes of RIBEYE and palm-Bassoon puncta from HEK293 cells expressing RIBEYE and palm-Bassoon (n = 20 cells, quantifications from 5 sample transfections). Volumes of synaptic ribbons from rat IHCs (n = 29 cells, 3 rats) have been plotted for comparison. E Box plot depicting data from **(D)**. Volumes of RIBEYE and palm-Bassoon puncta are comparable to each other (*P* > 0.99, Kruskal-Wallis test with *post hoc* Dunn’s multi-comparison), but on an average much smaller and considerably more variable when compared to volumes of RIBEYE and Bassoon puncta from rat IHCs respectively (*\*\*\*\*P* < 0.0001, Kruskal-Wallis test with *post hoc* Dunn’s multi-comparison). Overlaid plus signs represent individual spots, crosses represent mean values, central band indicates the median, whiskers represent 90/10 percentiles and boxes represent 75/25 percentiles. F RIBEYE puncta in HEK cells expressing RIBEYE and palm-Bassoon (n_spots_ = 864, N = 20 cells) appear more spherical than Bassoon puncta in the same cells (n_spots_ = 961, N = 20 cells; *\*\*\*\*P* < 0.0001, Kruskal-Wallis test with *post hoc* Dunn’s multiple comparison test). Note the similar trend in IHC synaptic ribbons where RIBEYE puncta are more spherical than Bassoon puncta (n_spots_ = 290 for RIBEYE and n_spots_ = 314 for Bassoon, N = 29 cells, 3 rats; *\*\*\*\*P* < 0.0001, Kruskal-Wallis test with *post hoc* Dunn’s multiple comparison test). Overlaid plus signs represent individual spots, crosses represent mean values, central band indicates the median, whiskers represent 90/10 percentiles and boxes represent 75/25 percentiles.

We compared the RIBEYE and Bassoon signal intensities at the periphery *versus* inside of the cell in randomly selected single sections from confocal stacks of HEK cells expressing RIBEYE and palm-Bassoon (N = 9 cells) and HEK cells expressing RIBEYE and full-length Bassoon (N=9 cells). We found an increased peripheral distribution of both RIBEYE and Bassoon when using palm-Bassoon (Fig. 2C, *\*\*\*\*P* < 0.0001, Mann-Whitney-Wilcoxon test), implying successful membrane targeting of RIBEYE by palm-Bassoon. In each transfection, ∼10% cells showed co-expression of both RIBEYE and palm-Bassoon (for representative sample overview see Supplementary Fig. 2). Of those, cells expressing discrete RIBEYE-palm Bassoon structures were discernible by the characteristic RIBEYE distribution along the periphery of the cell. This peripheral distribution in turn seemingly depends upon RIBEYE and palm-Bassoon expression ratios (shown in Supplementary Fig. 2B) as cells with little to no palm-Bassoon expression show predominantly cytosolic RIBEYE puncta and were not used for analysis. For simplicity, we henceforth refer to the structures composed of RIBEYE and palm-Bassoon in HEK293 cells as *SyRibbons* (for *synthetic* ribbons).

We next performed three-dimensional surface renderings using *Imaris* 9.6 (Oxford Instruments) to assess these structures. In a given cell, only structures with colocalizing RIBEYE and palm-Bassoon immunofluorescence were considered for analysis to exclude occasional non-membrane localized spots (on average 51.75 ± 40.84 RIBEYE surfaces colocalizing with Bassoon surfaces per cell; N = 20 cells). The volumes of RIBEYE and palm-Bassoon surfaces of *SyRibbons* were smaller on average and more variable (average volume ± standard deviation (SD) = 0.19 ± 0.23 µm^3^ with a coefficient of variation (CV) = 1.23 for RIBEYE; n_spots_ = 864 and volume = 0.17 ± 0.18 µm^3^ with CV = 1.10 for Bassoon; n_spots_ = 951; data from N = 20 cells, quantifications from 5 sample transfections) than volumes of synaptic ribbons and Bassoon immunofluorescent puncta from rat IHCs (volume = 0.29 ± 0.17 µm^3^, CV = 0.59 for RIBEYE; n_spots_ = 290, and volume = 0.27 ± 0.18 µm^3^, CV = 0.67 for Bassoon; n_spots_ = 314, data from N = 29 cells), (Fig. 2D, E). Moreover, volumes of RIBEYE surfaces show a high positive correlation to volumes of corresponding palm-Bassoon surfaces (*P_r_* = 0.778, *\*\*\*\*P* < 0.0001), implying that larger palm-Bassoon structures may recruit bigger RIBEYE structures to the plasma membrane. We also note that the volume of RIBEYE surfaces in *SyRibbons* appears smaller and well-regulated in contrast to the predominantly large, cytosolic RIBEYE assemblies in cells co-expressing RIBEYE and full-length Bassoon (volume = 0.30 ± 0.36 µm^3^, n_spots_ = 460, N = 10 cells, *\*\*\*\*P* < 0.001, Mann-Whitney-Wilcoxon Test). In turn, Bassoon clusters seemed regulated by co-expressed RIBEYE: Bassoon surfaces at the plasma membrane were larger at *SyRibbons* than in the absence of RIBEYE in cells only expressing palm-Bassoon (volume = 0.14 ± 0.21 µm^3^, n_spots_ = 2217, N = 13 cells, *\*\*\*\*P* < 0.001, Mann-Whitney-Wilcoxon Test). RIBEYE clusters constituting *SyRibbons* appeared more spherical than the plate-like Bassoon structures in the same cells (sphericity Ψ = 0.91 ± 0.05, n_spots_ = 864 for RIBEYE versus Ψ = 0.86 ± 0.07, n_spots_= 951 for Bassoon, data from N = 20 cells; *\*\*\*\*P* < 0.0001, Kruskal-Wallis test with *post hoc* Dunn’s multiple comparison test). This follows the same trend as in rat IHCs where RIBEYE spots indeed appear more spherical (Ψ = 0.90 ± 0.05, n_spots_ = 290) compared to Bassoon spots (Ψ = 0.76 ± 0.10, n_spots_ = 314); *\*\*\*\*P* < 0.0001, Kruskal-Wallis test with *post hoc* Dunn’s multiple comparison test, data from N = 29 cells (Fig. 2F). Nonetheless, the fact that next to structures with volumes comparable to IHC ribbons, we also encountered smaller and larger structures, likely reflects poorer regulation of RIBEYE and palm-Bassoon expression in the heterologous system.

Next, we analyzed *SyRibbons* in situ using cryo-electron tomography (cryo-ET), which capitalizes on cell vitrification to obtain near-native preservation. Plunge-frozen HEK cells transfected with RIBEYE-GFP and untagged palm-Bassoon were vitrified and subsequently milled using a cryo-focused ion beam (cryo-FIB; Supplementary Fig. 3A) to produce 150 nm-thick lamellae as previously described (Rigort *et al*, 2010; Pierson *et al*, 2024, see methods). Using a light microscope integrated in the cryo-FIB chamber, we targeted cell areas showing peripheral GFP fluorescence corresponding to *SyRibbons* (Supplementary Fig. 3B). Lamellae were subsequently transferred to a cryo-transmission electron microscope, and tomographic tilt-series were acquired at fluorescence locations. The tomograms showed that GFP-positive RIBEYE spots corresponded to electron-dense structures (Fig 3A-B), as is characteristic of the synaptic ribbon (Robertis & Franchi, 1956; Smith & Sjöstrand, 1961). These electron-dense *SyRibbons* appeared to be approx. 300 – 800 nm in size and were ovoid or ellipsoidal in shape (Fig 3B-C, Supplementary Fig. 3C). Some *SyRibbons* displayed a hollow core (Supplementary Fig. 3Ci and 3C_iii_) and on closer inspection, some appeared to have a multi-lamellar ultrastructure (Supplementary Fig. 3Cii), both of which have been reported previously for natively expressed synaptic ribbons in the inner ear and the retina (Michanski *et al*, 2023, 2019; Sobkowicz *et al*, 1982; Liberman, 1980; Stamataki *et al*, 2006; Wichmann & Moser, 2015). We captured a *SyRibbon* positioned within 100 nm from the plasma membrane (Fig 3C). Although bona fide ribbons display a halo of SVs tethered on their surface, we did not observe any obvious accumulation of vesicular structures around these *SyRibbons*, which is not unexpected for synapse-naïve HEK cells. This is in contrast to a previous report in R28 retinal progenitor cells, where heterologously expressed RIBEYE was shown to recruit vesicles (Magupalli *et al*, 2008). Intriguingly, in some tomograms, we observed membrane-bound *SyRibbons* (Supplementary Fig. 3Ci_-iv_), which may indicate an autophagic engulfment of some of these overexpressed structures.

**Figure 3.**
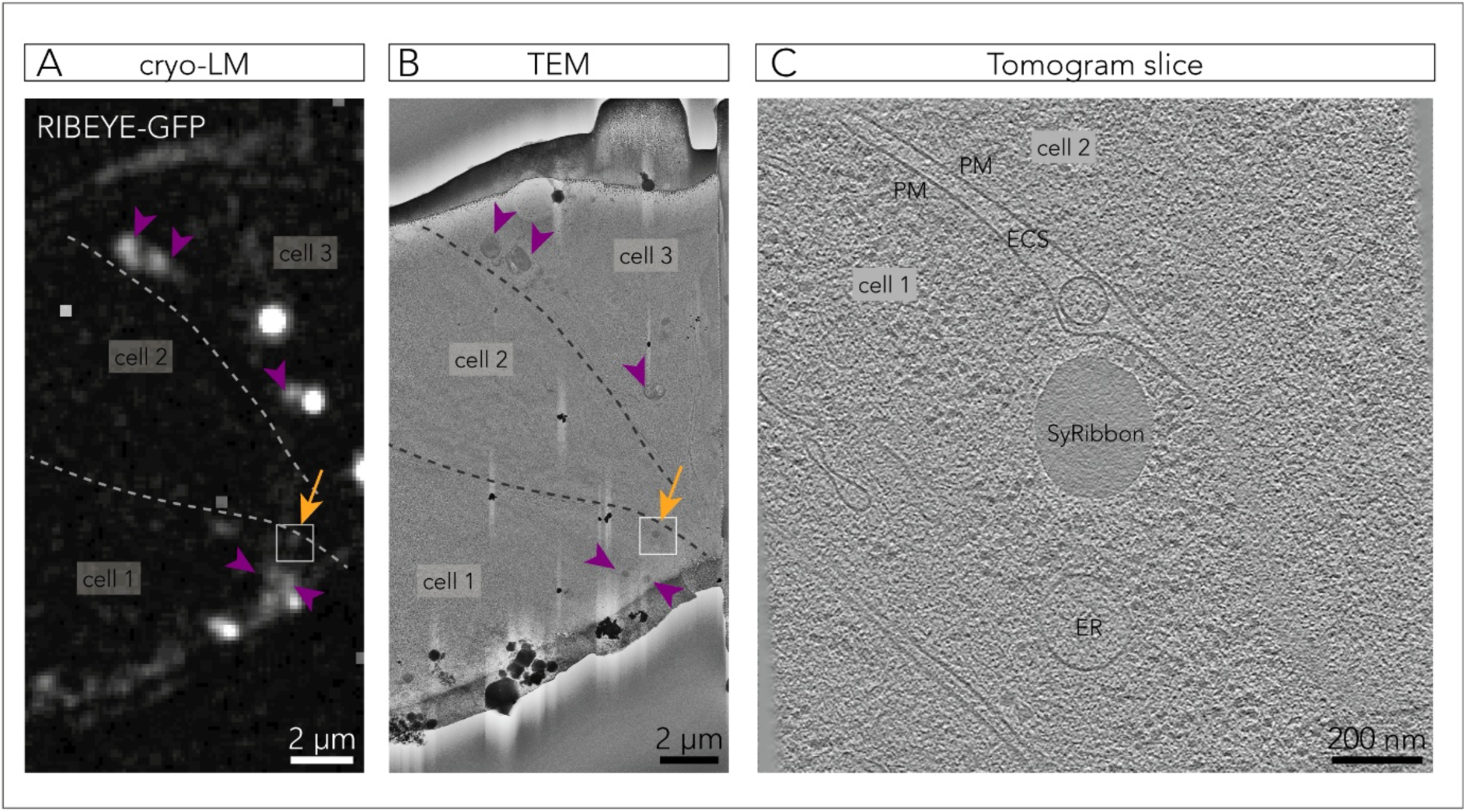
Cryo-correlative microscopy captures membrane-localised *SyRibbons*. **A**. RIBEYE-GFP signal on a 150 nm thick lamella, revealed by fluorescent light microscopy within the cryo-FIB chamber (cryo-LM). Dotted lines delineate cell membranes. The orange arrow indicates a plasma membrane-proximal GFP fluorescence, whereas purple arrow heads point to cytosolic RIBEYE aggregates. The tomogram shown in (C) was acquired at the boxed region. The image underwent background subtraction for better visualization. **B**. Transmission electron microscopy (TEM) image of the lamella in A. Orange arrow and purple arrowheads locate the respective regions in (A), showing that GFP-positive spots correlated to electron dense bodies. Dotted lines delineate plasma membranes. The tomogram shown in (C) was taken at the boxed region. **C**. Tomogram slice showing a *SyRibbon* acquired at the boxed region in (B). *PM*: plasma membrane, *ECS*: extracellular space *ER*: endoplasmic reticulum.

### RBP2 tethers Ca_V_1.3 channels

To study the potential of *SyRibbons* to cluster Ca^2+^ channels, we employed HEK293 cells stably expressing an inducible transgene of Ca_V_1.3α_1_ along with constitutive transgenes for Ca_V_β_3_ and Ca_V_α_2_8_1_. We either used tetracycline for inducible Ca_V_1.3α_1_ channel expression (Fig. 4A) or performed transient transfections with Ca_V_1.3 constructs containing an N-terminal EGFP- (Fig. 4B) or Halo-tag (Fig. 4C) for direct fluorescence imaging. Expression of either Ca^2+^ channel complex resulted in small clusters at the plasma membrane as represented in Fig. 4D (average volume 0.08 ± 0.13 µm^3^, n_spots_ = 949, N = 12 cells for untagged inducible Ca_V_1.3α_1_; 0.07 ± 0.08 µm^3^, n_spots_ = 392, N = 11 cells for EGFP-Ca_V_1.3; and 0.07 ± 0.09 µm^3^, n_spots_ = 1281, N = 16 cells for Halo-Ca_V_1.3α; *P_untagged/EGFP_* = 0.547, *P_untagged/Halo_* > 0.999, *P_Halo/EGFP_* = 0.582, Kruskal-Wallis test with *post hoc* Dunn’s multiple comparison test).

**Figure 4.**
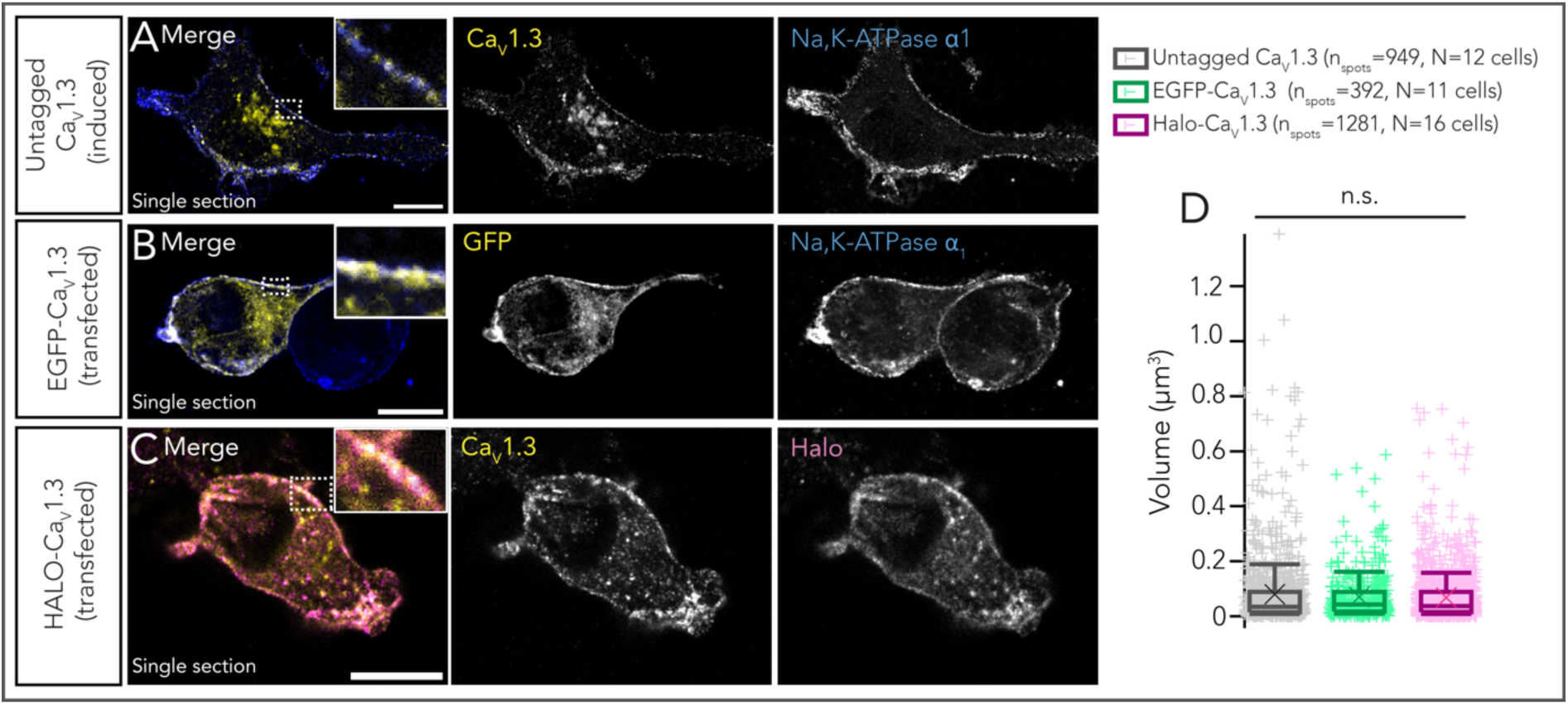
HEK293 cells expressing CaV1.3. A Representative confocal section of a HEK293 cell stably expressing an inducible Ca_V_1.3α transgene (untagged) along with constitutive transgenes for Ca_V_β_3_ and Ca_V_α_2_8_1._ Immunostainings have been performed using antibodies against Ca_V_1.3 (green) and Na, K-ATPase α_1_ (blue, labeling the plasma membrane). Scale bar = 10 µm. B Representative confocal section of a HEK293 cell stably expressing Ca_V_β_3_ and Ca_V_α_2_8_1,_ and transiently transfected with an N-terminal EGFP-tagged Ca_V_1.3 construct. Immunostainings have been performed using antibodies against GFP (green) and Na, K-ATPase α_1_ (blue). Scale bar = 10 µm. C Representative confocal section of a HEK293 cell stably expressing Ca_V_β_3_ and Ca_V_α_2_8_1_ and transiently transfected with an N-terminal Halo-tagged Ca_V_1.3 construct. Immunostainings have been performed using antibodies against Ca_V_1.3 (green) and Halo tag (magenta). Scale bar = 10 µm. D Expression of either of the Ca^2+^ channel complexes result in clusters of comparable sizes, (*P* > 0.05, Kruskal-Wallis test with *post hoc* Dunn’s multiple comparison test). Overlaid plus signs represent individual spots, crosses represent mean values, central band indicates the median, whiskers represent 90/10 percentiles and boxes represent 75/25 percentiles.

Co-expression of palm-Bassoon did not show colocalization with Ca_V_1.3, which is in line with observations with full-length Bassoon (Frank *et al*, 2010) and the same applied when co-expressing RIBEYE and Ca_V_1.3, arguing against a direct interaction for both (Supplementary Fig. 4). Rab-interacting molecule-binding protein (RIM-BP or RBP) links Bassoon to Ca^2+^ channels (Davydova *et al*, 2014) and hence was an interesting candidate for tethering Ca_V_1.3 to Bassoon clusters. RBP has been shown to interact with Ca_V_1.3 (Hibino *et al*, 2002) and to be required for normal Ca_V_1.3 Ca^2+^ channel clustering at the IHC ribbon synapse (Krinner *et al*, 2017, 2021). Indeed, when co-expressing RBP2 with palm-Bassoon and Ca_V_1.3, we found juxtaposed palm-Bassoon and Ca_V_1.3 both with immunolabeled (Fig. 5A) and live-labeled (Fig. 5B, C) proteins indicating successful clustering of Ca_V_1.3 at *synthetic* AZs.

**Figure 5.**
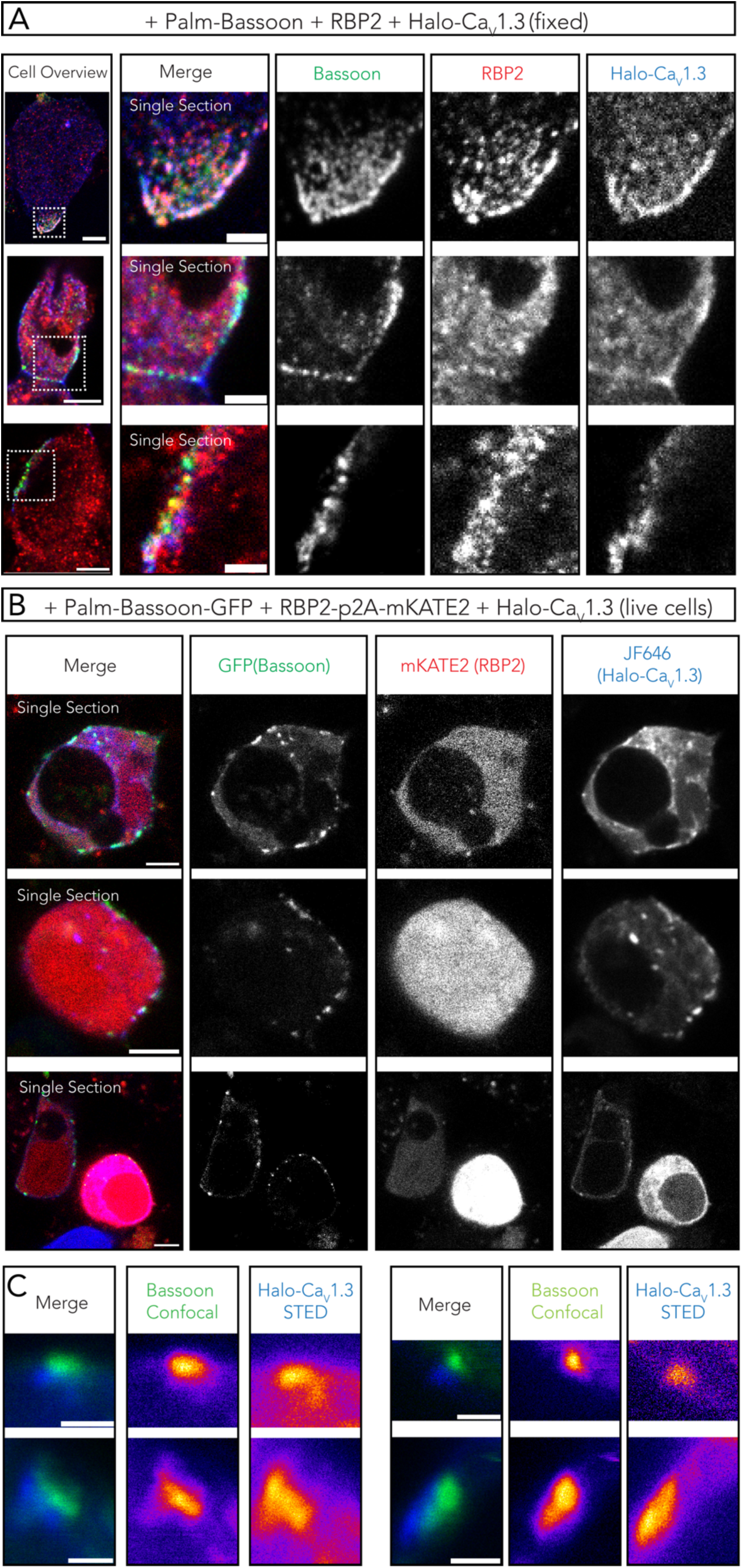
RBP2 bridges palm-Bassoon and Ca_V_1.3 to form supra-molecular assemblies at the plasma membrane. A Representative confocal images (single sections) of HEK293 cells (fixed) transfected with palm-Bassoon (green), RBP2 (red) and N-terminal Halo-tagged Ca_V_1.3 (blue). The three colocalizing proteins appear to form supra-molecular assemblies at the cell membrane. Scale bar = 5 µm for left panel showing cell overviews, 2 µm for right panels showing zoom-ins. B Representative confocal images (single sections) of live HEK293 cells transfected with palm-Bassoon-GFP (green), RBP2-p2A-mKATE2 (red, representing only mKATE2 signal and not RBP2 localization) and Halo-Ca_V_1.3 (blue, labeled by with JF-646 HaloTag Ligand before imaging). Scale bar = 5 µm. C Exemplary super-resolution STED images from live-labeled samples in (B) showing juxtaposition of palm-Bassoon and Halo-Ca_V_1.3 signal at the plasma membrane in RBP2 positive cells. Scale bar = 1 µm.

We next performed ruptured whole-cell patch-clamp to assess if RBP2 and/or palm-Bassoon or RIBEYE co-expression results in changes in Ca_V_1.3 Ca^2+^ currents. After tetracycline treatment for Ca_V_1.3α_1_ expression (18-24 hours), cells were transfected with both RBP2-p2A-mKATE2 and palm-Bassoon-GFP, with only palm-Bassoon-GFP, only RBP2-p2A-mKATE2 or RIBEYE-GFP constructs (Supplementary Fig. 5A). After 18-24 hours, the cell culture media was changed, and cells were recorded 24-36 hours afterwards. We used 10 mM [Ca^2+^]_e_ and recorded Ca^2+^ current (density) - voltage relations (IVs) ∼1 minute after establishing the whole-cell configuration by applying step depolarizations of 20 ms from −86.2 mV to 58.8 mV in 5 mV increments. Ca^2+^ current density appeared to be augmented upon expression of both RBP2 and palm-Bassoon when compared with induced-only controls (Supplementary Fig. 5B, D; *\*\*P* = 0.0041, Kruskal-Wallis test with *post hoc* Dunn’s multiple comparison test). However, Ca^2+^ current density in cells expressing only RBP2 or Palm-Bassoon did not significantly differ from either induced-only controls or cells expressing both proteins (*P* > 0.05, Kruskal-Wallis test with *post hoc* Dunn’s multiple comparison test). For cells expressing only RIBEYE, we also did not observe any changes (*P* > 0.999, Kruskal-Wallis test with *post hoc* Dunn’s multiple comparison test). We did not observe any noticeable differences in voltage-dependence, activation kinetics or inactivation kinetics of Ca_V_1.3 Ca^2+^ current upon expression of any of these AZ proteins (Supplementary Fig. 5C, E, F; Supplementary Fig. 6A, B).

### Larger synthetic ribbon-type AZs establish larger Ca_V_1.3 Ca^2+^ channel clusters

Next, we tetra-transfected HEK293 cells with palm-Bassoon, Halo-tagged Ca_V_1.3α_1_, RBP2 and RIBEYE to explore the impact of co-expressing of RIBEYE on the clustering of Ca_V_1.3 Ca^2+^ channels (Fig. 6A for transfection scheme). Using confocal imaging of immunofluorescently labelled RIBEYE, Halo-CaV1.3 and Bassoon (RBP2 immunofluorescence was imaged using epifluorescence to select tetra-transfected HEK293 cells), we observed colocalization of Ca_V_1.3α1, Bassoon and RIBEYE (Fig. 6B). Line profiles drawn tangentially along the plasma membrane showed that beyond the general distribution of a Ca_V_1.3α_1_ signal in the plasma membrane, Ca_V_1.3α_1_ signal hotspots occurred underneath the *SyRibbons* (Fig. 6C). The signal intensity of RIBEYE immunofluorescence positively correlated with the Ca_V_1.3 signal immunofluorescence, (*P_r_* = 0.839 and 0.886 from two sample line scans, *\*\*\*\*P* < 0.0001), which has previously been reported for immunolabeled native ribbon-type AZs of mouse IHCs (Ohn *et al*, 2016). We further performed a pixel-based correlation analysis (Supplementary Fig. 7A, B) which indicated a moderate to strong colocalisation of RIBEYE and Ca_V_1.3 upon co-expression of palm-Bassoon and RBP2. In contrast, a poor colocalization was found in cells expressing only RIBEYE and Ca_V_1.3; average Pearson’s colocalisation coefficient = 0.41 versus 0.24; average Mander’s overlap coefficient = 0.63 versus 0.34, N = 12 and 8 cells respectively (*\*\*P* < 0.01m Wilcoxon’s Rank Test). Colocalisation quantifications for other transfection combinations shown in Fig. 2, 5 and Supplementary Fig. 4 have been collectively depicted in Supplementary Fig. 7. We next turned to 2-color STED imaging which provided improved spatial resolution and highlighted the confined localization of Ca_V_1.3 Ca^2+^ -channel clusters underneath *SyRibbons* as shown in Fig. 6D.

**Figure 6.**
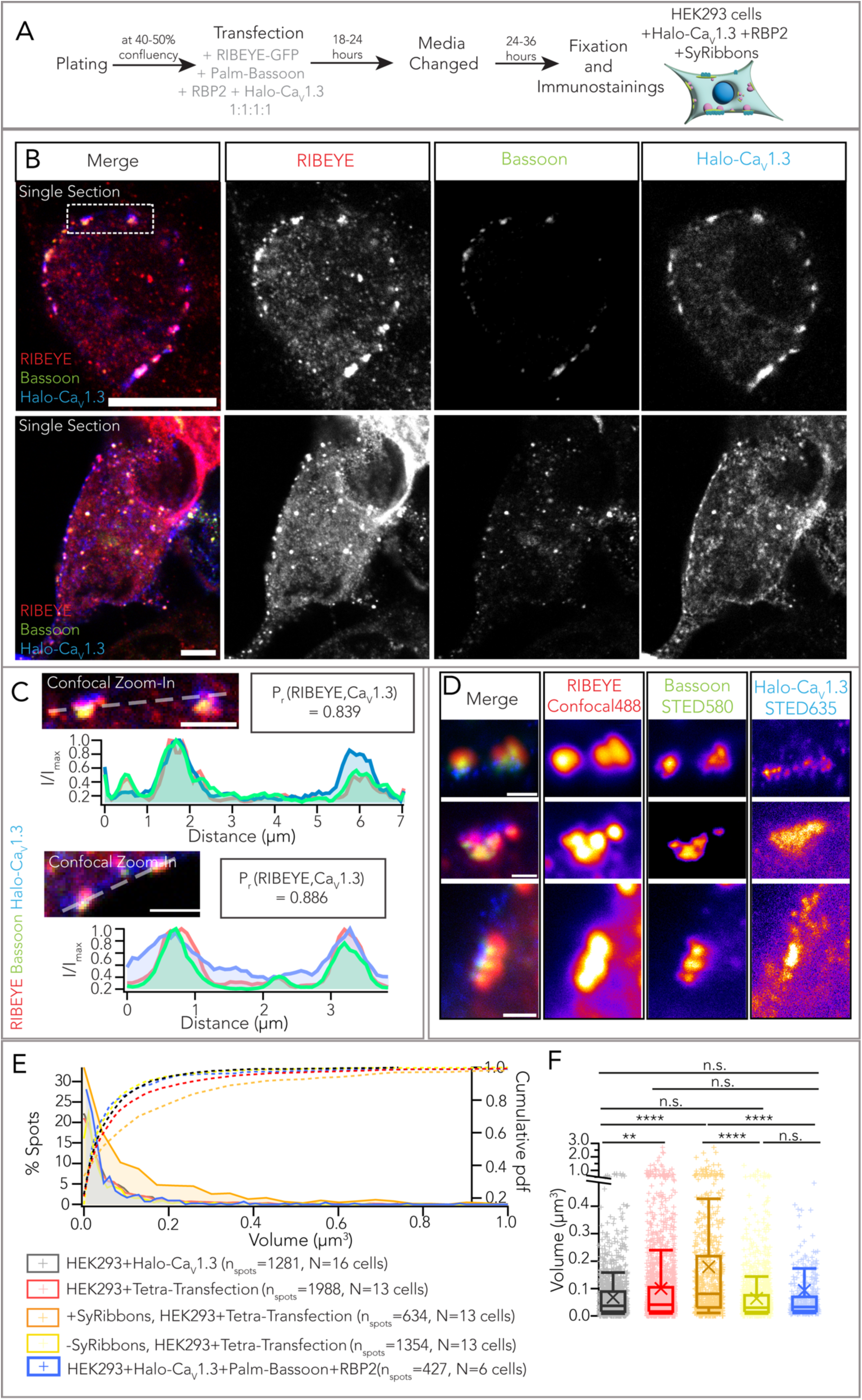
Synthetic ribbon-type active zones recruit Ca_V_1.3 Ca^2+^ channels. A Experimental scheme for expression of Halo tagged Ca_V_1.3, palm-Bassoon, RIBEYE-GFP and RBP2 in HEK293 cells. B Representative confocal images (single sections) of HEK293 cells transfected with RBP2 (not shown, expression confirmed using epifluorescence), RIBEYE-GFP (red), palm-Bassoon (green), Halo-Ca_V_1.3 (blue) shows colocalization of the three latter proteins at the plasma membrane. Scale bar = 5 µm. C Confocal zoom-ins from **(B)**. Line scans depict an increased Ca_V_1.3 signal intensity (blue) at sites where RIBEYE (red) and palm-Bassoon (green) clusters localize. Pearson’s correlation coefficients (*Pr*) were calculated along the line profiles for RIBEYE and Ca_V_1.3 signal intensity and indicate a high degree of correlation (∼0.8) between the spatial localization of the two. Scale bar = 2 µm. D Super-resolution STED images showing Ca_V_1.3 clusters localizing at the base of the *SyRibbons*. Scale bar = 500nm. E Distribution of volumes from confocal images of Ca_V_1.3 clusters colocalizing with *SyRibbons* (orange, n_spots_ = 634) and not colocalizing with *SyRibbons* (yellow, nspots = 1354) from HEK293 cells expressing Ca_V_1.3, RBP2 and *SyRibbons* (N = 13 cells). Pooled volumes of all Ca_V_1.3 clusters (with and without *SyRibbons*) from these cells have been shown in red (n_spots_ = 1988). Volumes of Ca_V_1.3 clusters from HEK293 cells expressing only Halo-Ca_V_1.3 (black, n_spots_ = 1281, N = 16 cells) and from cells expressing Halo-CaV1.3, RBP2 and palm-Bassoon (blue, n_spots_ = 427, N = 6 cells) have been plotted for comparison. F Box plot depicting data from (E). Ca_V_1.3 clusters appear larger on an average in cells expressing Ca_V_1.3, RBP2 and *SyRibbons versus* in cells expressing only Ca_V_1.3, (*\*\*P* = 0.0044). Within cells expressing all four proteins, Ca_V_1.3 clusters are much larger when colocalizing with *SyRibbons* versus when they do not (*\*\*\*\*P < 0.0001*). Ca_V_1.3 clusters colocalising with *SyRibbons* were in fact also larger than clusters in cells expressing only Ca_V_1.3 and cells expressing only RBP2 and palm-Bassoon (*\*\*\*\*P < 0.0001*). Statistical test: Kruskal-Wallis test with *post hoc* Dunn’s multiple comparison correction. Overlaid plus signs represent individual spots, crosses represent mean values, central band indicates the median, whiskers represent 90/10 percentiles and boxes represent 75/25 percentiles.

We performed surface renderings of confocally imaged Halo-tagged Ca_V_1.3α_1_, RIBEYE and palm-Bassoon immunofluorescence spots as described above for estimating the volume of *SyRibbons* and the Ca_V_1.3 Ca^2+^ channel clusters (Fig. 6E, F). This revealed significantly larger Ca_V_1.3 Ca^2+^ channel clusters when colocalizing with *SyRibbons* in HEK293 + Halo-Ca_V_1.3 + RBP2 + *SyRibbons* cells (average volume ± SD = 0.18 ± 0.28 µm^3^, n_spots_= 634, N = 13 cells, data representative of 5 transfections) compared to ribbonless (i.e. non-colocalized) Ca_V_1.3 Ca^2+^ channel clusters from the same cells (volume = 0.06 ± 0.10 µm^3^, n_spots_= 1354, N = 13 cells; *\*\*\*\*P* < 0.0001, Kruskal-Wallis test with *post hoc* Dunn’s multiple comparison test) and Ca_V_1.3 Ca^2+^ clusters in HEK293 cells solely expressing Halo-Ca_V_1.3 Ca^2+^ channels (volume = 0.07 ± 0.09µm^3^, n_spots_= 1281, N = 16 cells, data representative of 4 transfections; *\*\*\*\*P* < 0.0001, Kruskal-Wallis test with *post hoc* Dunn’s multiple comparison test). Ribbonless Ca_V_1.3 Ca^2+^ channel clusters did not differ significantly regardless if co-expressing RIBEYE (*P* = 0.099, Kruskal-Wallis test with *post hoc* Dunn’s multiple comparison test, see Fig. 6F). Moreover, volumes of Ca_V_1.3 cluster surfaces show a moderate positive correlation with volumes of corresponding RIBEYE surfaces (*P_r_* = 0.385, *\*\*\*\*P* < 0.0001), similar to previous observations at IHC ribbon synapses (Frank *et al*, 2009; Michanski *et al*, 2023; Ohn *et al*, 2016).

### Ca^2+^ imaging reveals a partial spatial confinement of Ca^2+^ signal at synthetic ribbon-type active zones

Finally, we set-out to functionally characterize synthetic ribbon-type AZs. We combined patch-clamp recordings of Ca_V_1.3 Ca^2+^ influx with spinning-disk confocal microscopy to visualize Ca^2+^ signals at the synthetic ribbon-type AZs in HEK293 cells. Following tetracycline-induction for Ca_V_1.3 expression, HEK293 cells were co-transfected with constructs expressing RIBEYE-GFP, untagged palm-Bassoon and RBP2-p2A-mKATE2, (Fig. 7A). We recorded mKATE2 positive cells that expressed peripheral GFP puncta, indicative of RIBEYE and palm-Bassoon co-expression. We employed peripheral GFP expression as a proxy of *SyRibbons* for functional analysis (Fig. 7B). Ruptured patch-clamp recordings were performed with 10 mM [Ca^2+^]_e_ and 10 mM intracellular EGTA to enhance the signal to background ratio for visualizing Ca^2+^ influx using the low-affinity red-shifted Ca^2+^ indicator Calbryte590 (100 µM, k_d_ = 1.4 µM). After loading the cell for approximately a minute, we first applied step depolarizations of 20 ms from −83 to +62 mV in 5-mV step increments to measure Ca^2+^ current (density)–voltage relations. We did not observe any changes in either the Ca^2+^ current density (*P* = 0.293, t-test) or the voltage-dependence of Ca^2+^ current (*P* = 0.780, t-test when comparing V_half_) when comparing cells expressing Ca_V_1.3 + RBP2 + *SyRibbons* with cells expressing only CaV1.3 (Fig. 7C and 7D). Next, we applied a depolarizing pulse to +2 mV for 500 ms and acquired images at a frame rate of 20 Hz (Fig. 7E). The intracellular Ca^2+^ signals mediated by Ca_V_1.3 Ca^2+^ influx were visualized as an increase in Calbryte590 fluorescence. The increase of Calbryte590 fluorescence appeared to spread throughout the HEK293 cell membrane with discrete regions of higher signal intensity at *SyRibbons* (exemplary cell shown in Fig. 7F with zoom-in at site with *SyRibbon* shown in 7G). We note that we did not find noticeable red fluorescence at *SyRibbons* in the absence of depolarization, which is similar to findings with Fluo-4FF in IHCs (Frank *et al*, 2009), but different from the situation for the large spherical presynaptic bodies in bullfrog hair cells that are stained by Fluo-3 (Issa & Hudspeth, 1996). Line profiles drawn tangentially to the membrane in composite ΔF image of Calbryte590 signal and RIBEYE-GFP showed a high correlation between the localization of *SyRibbons* and peak intensity of Calbryte590 fluorescence increase, indicating preferential Ca^2+^ signaling at the *SyRibbons* (two representative line scans shown in Fig. 7H, Pearson correlation coefficients of 0.783 and 0.757). Recordings were made from 24 cells (7 transfections) and ROIs (diameter = 2 µm) with and without *SyRibbons* were analyzed as shown in Fig. 7I. Calbryte590 ΔF_max_/F_0_ values were calculated for each ROI which on an average showed higher ΔF_max_/F_0_ values for ROIs with *SyRibbons* (on average ∼26% higher) than for ROIs without *SyRibbons* (*\*P* = 0.018, paired t-test, Fig. 7J). We note that occasionally in some cells we observed a higher Calbryte590 signal increase at sites without *SyRibbons*, and we attribute this to the likely presence of AZ-like clusters composed of RBP2 and palm-Bassoon in these regions which we could not visualize.

**Figure 7.**
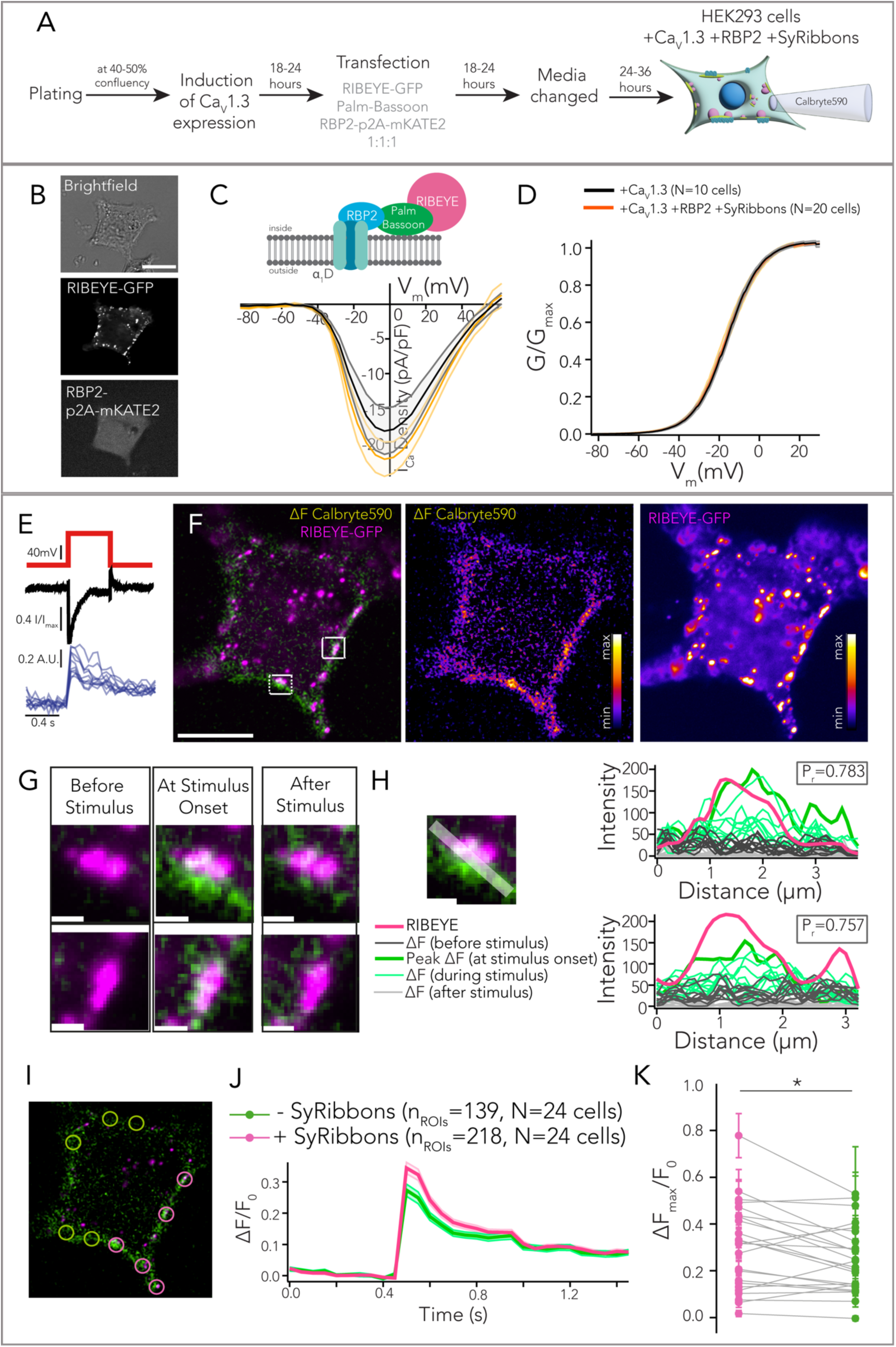
Ca^2+^ imaging reveals higher Ca^2+^ signal intensity underneath *SyRibbons*. A Experimental scheme for expression of *SyRibbons*, RBP2 and Ca_V_1.3 in HEK293 cells for patch clamp in combination with Ca^2+^ imaging using the low affinity Ca^2+^ indicator Calbryte590. B Exemplary cell used for Ca^2+^ imaging. Cells were identified by peripheral RIBEYE-GFP puncta as shown (indicative of RIBEYE and palm-Bassoon co-expression) and mKATE2 signal (indicative of RBP2 expression). C Current density-voltage (IV) relations from whole-cell patch clamp recordings of HEK293 cells expressing Ca_V_1.3 + RBP2 + *SyRibbons* (n=20, in orange) and only Ca_V_1.3 (n=10, in black). Lines represent mean current traces, shaded area represents ± SEM. Whole-cell Ca^2+^ current density does not appear to change upon co-expression of RBP2 and *SyRibbons* (*P* = 0.293, t-test). D The voltage-dependence of Ca^2+^ current influx does not seem to be altered upon co-expression of RBP2 and *SyRibbons* at the whole-cell level. E A depolarizing pulse to +2mV was applied to the cells for 500 ms and the increase in Calbryte590 fluorescence was measured by acquiring images at a frame rate of 20 Hz using a spinning disk confocal microscope. F Representative image of a HEK293 cell expressing *SyRibbons*, RBP2 and Ca_V_1.3 stimulated as described in (E). RIBEYE-GFP signal has been shown in magenta; the change in Calbryte590 fluorescence intensity (ΔF) upon Ca^2+^ binding (the frame at the onset of depolarization to +2mV) has been shown in green. Note the distinct Ca^2+^ signal “hotspots” that co-localize with *SyRibbons*. Scale bar = 10 µm. G Zoom-ins from (F). Frames before, at the onset of, and after stimulation have been shown. Note the localized Ca^2+^ influx at the base of the *SyRibbons*. Scale bar = 1 µm. H Line profiles drawn tangentially to the membrane in composite ΔF image of Calbryte590 signal and RIBEYE-GFP show a high correlation between the localization of *SyRibbons* and peak intensity of Calbryte590 fluorescence increase (*P_r_* = ∼0.7). Plots show intensity profiles along line scans for RIBEYE-GFP (magenta) and Calbryte590 (light grey for frames before stimulus, bold green for peak intensity at onset of stimulus, light green lines for decaying intensity during ongoing stimulus and dark grey for frames post-stimulus). I Regions of interest (ROIs) of 2 µm diameter were drawn at sites with (magenta circles) and without (green circles) *SyRibbons* and the corresponding Calbryte590 ΔF/F_0_ values were calculated for each ROI. J Plot of average (shaded area represents ± SEM) ΔF/F_0_ values from ROIs with and without *SyRibbons* (218 and 139 ROIs respectively from N = 24 cells, data from 7 transfections). K On average, Calbryte590 ΔF_max_/F_0_ was higher for ROIs with *SyRibbons* than ROIs without them in a given cell (*\*P* = 0.018, paired t-test). Dots represent mean of ΔF_max_/F_0_ from ROIs with (magenta) and without (green) *SyRibbons* averaged per cell, error bars represent ± SEM.

## Discussion

In this study, we reconstituted and characterized a minimal ribbon-type AZ model system using heterologous expression. Co-expressing Ca_V_1.3 Ca^2+^ channels with membrane targeted Bassoon for RIBEYE anchorage to the plasma membrane and attracting exogenous RBP2 for clustering Ca^2+^ channels in HEK293 cells led to structures recapitulating basic aspects of IHC ribbon synapses. Despite the poor regulation of transgene expression in the synapse-naïve HEK293 cells, the AZ-like structures, in the subset of cells were similar in morphology to the native IHC AZs. This applied to the *synthetic* ribbons as well as to the AZ-like clusters of Bassoon, RBP2 and Ca_V_1.3 Ca^2+^ channels. However, all three components exhibited variability beyond the substantial natural heterogeneity found among the IHC AZs (Moser *et al*, 2023). Although expression of RIBEYE does not seem to affect Ca_V_1.3 physiology and kinetics, we demonstrate enhanced clustering of Ca_V_1.3 channels and localization of Ca^2+^ influx at *SyRibbons*, with a mild increase in Ca^2+^ signal intensity. As *SyRibbons* partially resemble native hair cell AZs, we expect this easily available and experimentally accessible system to serve as an advanced testbed for functional interactions of AZ proteins and Ca^2+^ channels. Further efforts towards reconstituting SVs and their release sites will help to enhance the utility of these synthetic AZs.

### Synthetic active zone reconstitution model to study ribbon synapse assembly

By demonstrating that membrane-targeted Bassoon suffices to anchor ribbon-like RIBEYE assemblies, this work adds to the top-down and bottom-up evidence for a key role of Bassoon in attracting RIBEYE to the presynaptic density at the AZ in photoreceptors and hair cells (Dick *et al*, 2003; Khimich *et al*, 2005; Jing *et al*, 2013; tom Dieck *et al*, 2005). Additional expression of multi-domain proteins of the presynaptic density such as CAST (Ohtsuka *et al*, 2002; Inoue *et al*, 2006) might avoid the need for artificial palmitoylation of Bassoon for membrane targeting. Furthermore, our results support a model of a bidirectional control of AZ size and shape by RIBEYE and proteins of the presynaptic density. The amount of RIBEYE assembled in membrane localized *SyRibbons* appeared regulated strongly contrasting the seemingly unregulated cytosolic RIBEYE assemblies dominating HEK293 cells in the absence of membrane-targeted Bassoon. Vice versa, clusters of Bassoon at the plasma membrane were larger at *SyRibbons* than in the absence of RIBEYE. Finally, the size of *SyRibbons* seemingly scaled with the size of Ca_V_1.3 and Bassoon clusters similar to what is observed in IHCs (Ohn *et al*, 2016). Such a model is consistent with results from genetic perturbation of native ribbon synapses: i) disruption of RIBEYE leads to a disintegration of Bassoon into smaller subclusters in IHCs (Jean *et al*, 2018), and ii) disruption of CAST and ELKS reduces the size of the ribbon-type AZs in rod photoreceptors (tom Dieck *et al*, 2012; Hagiwara *et al*, 2018). We speculate that achieving the full extent of large ribbon-type AZs such as in rod photoreceptors as well as regulating the size of ribbon-type AZs at a specific synapse according to the precise functional demands requires such a functional interplay between proteins of the presynaptic density and RIBEYE. We note that other ribbon-resident proteins such as piccolino likely contribute to this fine-tuning of the size and shape of ribbon-type AZs, which will be an exciting question for future studies. Indeed, disruption of piccolino in IHCs reduced the average ribbon size (Müller *et al*, 2019; Michanski *et al*, 2023). One exciting example of ribbon synapse heterogeneity manifests itself in IHCs where synapses show a spatial size gradient that likely relates to the diverse molecular and functional properties of the postsynaptic SGNs (reviewed in Moser *et al*, 2023).

### Insights into presynaptic Ca^2+^ channel clustering and function

Current discussion of the clustering of Ca_V_ Ca^2+^ channels offers two different models: low affinity protein-protein interactions (e.g. Hibino *et al*, 2002; Kaeser *et al*, 2011) via specific domains and liquid-liquid phase separation involving intrinsically disordered domains (e.g. Heck *et al*, 2019). Work on ribbon synapses has considered layers of organizing Ca_V_ Ca^2+^ channels at the AZ that seem more compatible with the former model: i) “micro-clustering” to which the synaptic ribbon contributes as a super-scaffold (Jean *et al*, 2018; Neef *et al*, 2018; Frank *et al*, 2010; Maxeiner *et al*, 2016) and ii) tethering and “nano-clustering” by RBPs, RIMs, CAST and ELKS (Liu *et al*, 2011; Grabner *et al*, 2015; Jung *et al*, 2015; Krinner *et al*, 2017; Luo *et al*, 2017; Hagiwara *et al*, 2018). Moreover, there is converging evidence for nanoscale coupling of few Ca_V_ Ca^2+^ channels to SV release sites (Brandt, 2005; Jarsky *et al*, 2010; Wong *et al*, 2014; Maxeiner *et al*, 2016; Özçete & Moser, 2021; Grabner & Moser, 2021; Jaime Tobón & Moser, 2023) for which a role of the ribbon has been more controversial (e.g. Maxeiner *et al*, 2016; Grabner & Moser, 2021). MINFLUX nanoscopy of rod photoreceptor AZs recently revealed a well-ordered double-line array topography of Ca_V_ Ca^2+^ channels, RIM, Bassoon and ubMunc13-2 at the presynaptic membrane on both sides of the ribbon (Grabner *et al*, 2022).

Clearly, we could not fully reconstitute this complex organization with the minimal set of molecular players at *SyRibbons* in HEK293 cells. Yet, we demonstrate a positive effect of *SyRibbons* on local Ca^2+^ signaling, recapitulating microclustering of Ca_V_1.3 Ca^2+^ channels at IHC ribbon synapses. However, we note that the functional impact of ribbon-mediated clustering of Ca_V_1.3 Ca^2+^ channels remains unclear at the moment. Our Ca^2+^ imaging data seems more reminiscent of previous work on ribbon-type AZs of immature IHCs (Wong *et al*, 2014) where a broader distribution of Ca_V_1.3 and ribbons go along with an immature state of the IHC AZs. We also note that the difference in Ca^2+^ signalling near or apart *SyRibbons* is rather subtle. We propose two explanations in this regard: (i) the use of an overexpression system diminishes the contrast of the Ca^2+^ signal between AZ-like domains and the regular membrane. Overexpressed Ca_V_1.3 is likely distributed diffusely across the plasma membrane in these cells along with more localised Ca_V_1.3 clusters observed at *SyRibbons* (see Fig. 6B lower panel). (ii) While the presence of *SyRibbons* appears to promote the formation of larger Ca_V_1.3 channels and partially localise the Ca^2+^ signal, it is unlikely that *SyRibbons* recruit a larger Ca^2+^ channel complement. IHCs of RIBEYE KO mice were reported to show normal Ca^2+^ current amplitudes at the whole-cell and single-AZ level (Jean *et al*, 2018), while the spatial spread of the Ca^2+^ signal was more diffuse in the absence of the ribbon.

We propose that *SyRibbons* scaffold Ca_V_1.3 channel nano-clusters in a non-native heterologous expression system resulting in an increased channel density in regions with *SyRibbons*. In support of this hypothesis, the Ca^2+^ signal averaged at *SyRibbons* showed a mildly increased maximal amplitude when compared with diffuse Ca^2+^ channel distribution in the absence of *SyRibbons.* Instead, the total number of channels may be unaffected, which would explain why Ca^2+^ current amplitudes at the whole-cell level are not impacted by expression of *SyRibbons*. A clear limitation in the study therefore arises from the use of an overexpression system, making it necessary to interpret results with caution and requiring *in vivo* and *ex vivo* validation. Future use of polycistronic gene expression or stable cell lines expressing all components could help with more consistent expression of the components and make the system become even more widely applicable.

Another limitation of the synthetic AZ system in HEK293 cells is the lack of SVs and the molecular machinery required for SV release. Nonetheless, our work paves the way for future studies including stable co-expression of AZ proteins including of the release site marker Munc13-1 (Sakamoto *et al*, 2018; Böhme *et al*, 2016) and potential delivery of SV machinery via co-expression (Park *et al*, 2021). Alternatively, using neurosecretory cells such as pheochromocytoma cells (PC12) or adrenal chromaffin cells in primary culture, offer the advantage of providing synaptic-like microvesicles and a large set of components of the presynaptic machinery such that expression of exogenous proteins might be limited to RIBEYE. On the other hand, the complex molecular background involving various types of Ca_V_ channels will likely complicate the interpretation. Regardless, *SyRibbons* in cultured cells offer great availability and as adherent cells, are well accessible to sophisticated analysis by techniques such MINFLUX nanoscopy and cryo-electron tomography.

## Materials and Methods

### HEK cell culture and transfections

In this study, we used human embryonic kidney (HEK293) cells stably expressing a tetracycline-inducible human Ca_V_1.3 pore forming α1-D subunit transgene (CACNA1D, NM_000720.2), and showing constitutive expression of Ca_V_β3 (CACNB3, NM_000725.2) and Ca_V_α_2_8-1 (CACNA2D1, NM_000722.2). The stably expressing cell line was acquired from Charles River Laboratories, Cleveland, Ohio, USA (Cat. No. CT6232). The cells were cultured as per product protocols in Dulbecco’s Modified Eagle Medium (DMEM) containing GlutaMax, high glucose and pyruvate (Gibco, Life Tech., 31966047), supplemented with 10% Fetal Bovine Serum (FBS; Gibco, Life Tech., A5256701) and 100 units/ml of Penicillin-Streptomycin (Gibco, Life Tech., 15070063). Additionally, the media was supplemented with 0.6 µM of Isradipine (Sigma Aldrich, I6658) and the following selection antibiotics (in mg/ml): 0.005 Blasticidin (Invivogen, ant-bl-05), 0.50 Geneticin (G418 Sulfate; Gibco, Life Tech., 10131027), 0.10 Zeocin (InvivoGen, ant-zn-05) and 0.04 Hygromycin (Thermo Fisher, 10687010). Cells were cultured in a humidified incubator at 37°C with 5% (v/v) CO_2_ saturation. Cells were split at ∼70% confluency every 3-4 days to prevent adverse effects on cell growth and channel expression, by dissociating the cell using Accutase (Sigma-Aldrich, A6964). The passage number did not exceed more than 26 passages. For experiments, cells were grown in media lacking selection antibiotics. For induction of α_1_-D subunit expression, cells were treated with selection antibiotic-free medium containing 3 µg/ml tetracycline. For transfection, 6.8 µg polyethylenimine (PEI, 25kDa linear, Polysciences, 23966) was added along with a total of 2 µg of DNA to a final volume of 100 µl of DMEM (without FBS). The PEI/DNA mixture was thoroughly mixed and allowed to incubate for 30 min at room temperature before being added to the cells (30 µl/well in a 24-well-plate containing 500 µl media, cell confluency ∼50-70%). For all experiments with co-transfections, we used an equimolar ratio of DNA (total amount of DNA was kept 2 µg in a 100 µl transfection mix). After 18-24 hours, transfection media was replaced with fresh media devoid of selection antibiotics. Cells were used for experiments (immunocytochemistry, patch clamp and Ca^2+^ imaging) 24-48 hours after changing the media or as specified. If cell confluency was too high, cells were reseeded to an appropriate confluency and allowed to settle for at least 12 hours before commencement of experiments.

### Expression vectors for RIBEYE, Bassoon, Ca_V_1.3 and RBP2

The RIBEYE-GFP construct used consisted of a human RIBEYE cDNA cloned inframe into a pAAV-GFP vector driven by a hybrid CMV-enhancer-human-beta-actin promoter (CMV-HBA) and containing a downstream Woodchuck Hepatitis Virus (WHV) Posttranscriptional Regulatory Element (WPRE) for mRNA stabilization. All Bassoon constructs encode rat Bassoon. The untagged full-length constructs were cloned in the pCS2+ vector. For the generation of palm-Bassoon constructs, we used a pEGFP-C1 vector backbone with a CMV promoter and kanamycin resistance cassette. The insert comprised of a palmitoylation consensus sequence of GAP43 (ATGCTGTGCTGTATGAGAAGAACCAAACAGGTTGAAAAGAATGATGAGGACCAAAAGATTTCCGGACTCAGATCTCGAG), followed by a cDNA sequence encoding amino acids 95-3938 of rat Bassoon, and a C-terminal monomeric GFP. For the untagged version, the GFP tag was replaced by a STOP codon. To generate the Halo-Ca_V_1.3 plasmid (Schwenzer *et al*, 2024), human Ca_V_1.3 cDNA (accession number NM_001128840.2) was de-novo synthesized and assembled into a HaloTag vector (Promega G7721) using restriction cloning, thus encoding an N-terminal fusion of Ca_V_1.3 to HaloTag linked by a ‘GGS’ sequence. The analogous GFP-Ca_V_1.3 plasmid was generated by exchanging the HaloTag sequence for mEGFP using restriction cloning. The RBP2 construct with an mKATE2 tag comprised of an insert coding for mouse RBP2 (accession ID NP_001074857.1) with a p2A cleavage site followed by a C-terminal mKATE2 cloned inframe into a f(syn)w-mKATE2-p2A vector driven by a CMV promoter-enhancer and containing downstream WPRE sequence. The untagged version was made by cloning the RBP2 cDNA from this plasmid and replacing RIBEYE-GFP in the pAAV-CMV-HBA-RIBEYE-GFP-WPRE plasmid by in-fusion cloning (Takara Bio USA, Inc.).

### Immunocytochemistry and Imaging

HEK293 cells plated on poly-L-lysine coated coverslips were fixed with 99% chilled methanol at −20°C for 2 minutes as previously described (Picher *et al*, 2017b). The coverslips were washed thoroughly three times with PBS at room temperature (5 – 10 min). Blocking and permeabilization was performed with GSDB (goat serum dilution buffer: 16% normal goat serum, 450 mM NaCl, 0.3% Triton X-100, 20mM phosphate buffer, pH ∼7.4) for 45-60 minutes at room temperature. Samples were incubated with respective primary antibodies (diluted in GSDB, refer to Table 2) overnight at 4°C or for 2 hours at RT. The samples were then washed three times (5–10 minutes each) with wash buffer (450 mM NaCl, 0.3% Triton X-100, 20mM phosphate buffer, pH ∼ 7.4). Incubation with appropriate secondary antibodies (also diluted in GSDB refer to Table 2) was performed for 1 hour at room temperature in a light-protected wet chamber. Lastly, the coverslips were washed three times with wash buffer (5-10 minutes each) and one final time with PBS, before mounting onto glass slides with a drop of fluorescence mounting medium (Mowiol 4-88, Carl Roth). For live cell imaging of cells transfected with Halo-Ca_V_1.3 (Fig. 4B, C), cells plated on glass-bottomed dishes were treated with Janelia Fluor 646 HaloTag Ligand (Promega, Cat. No. GA1120) at a final labelling concentration of 200nM (in cell culture media). Cells were incubated with the ligand for ∼60 min at 37°C with 5% (v/v) CO_2_ saturation, after which the media was replaced with an equal volume of fresh warm culture media.

**Table 1:**
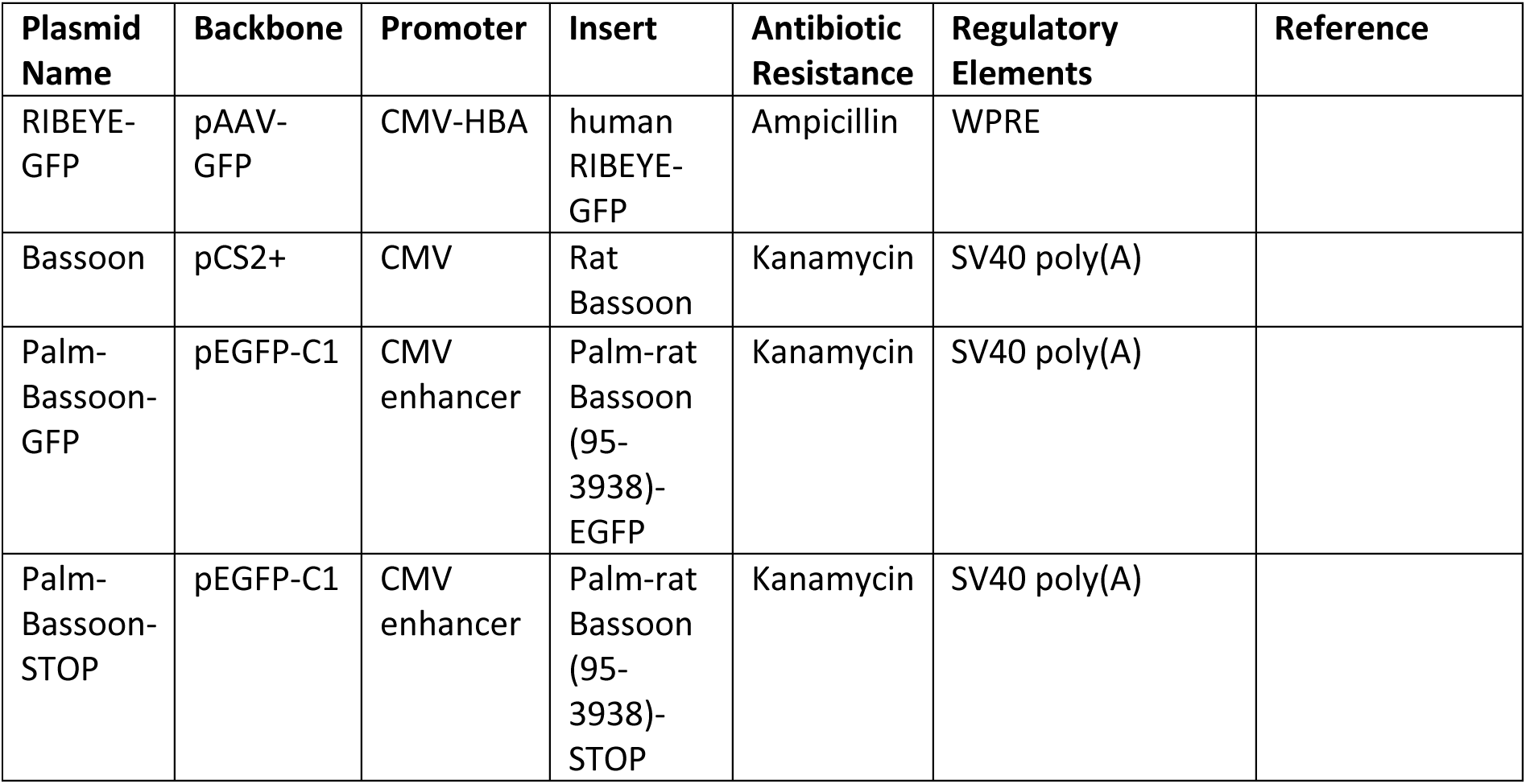
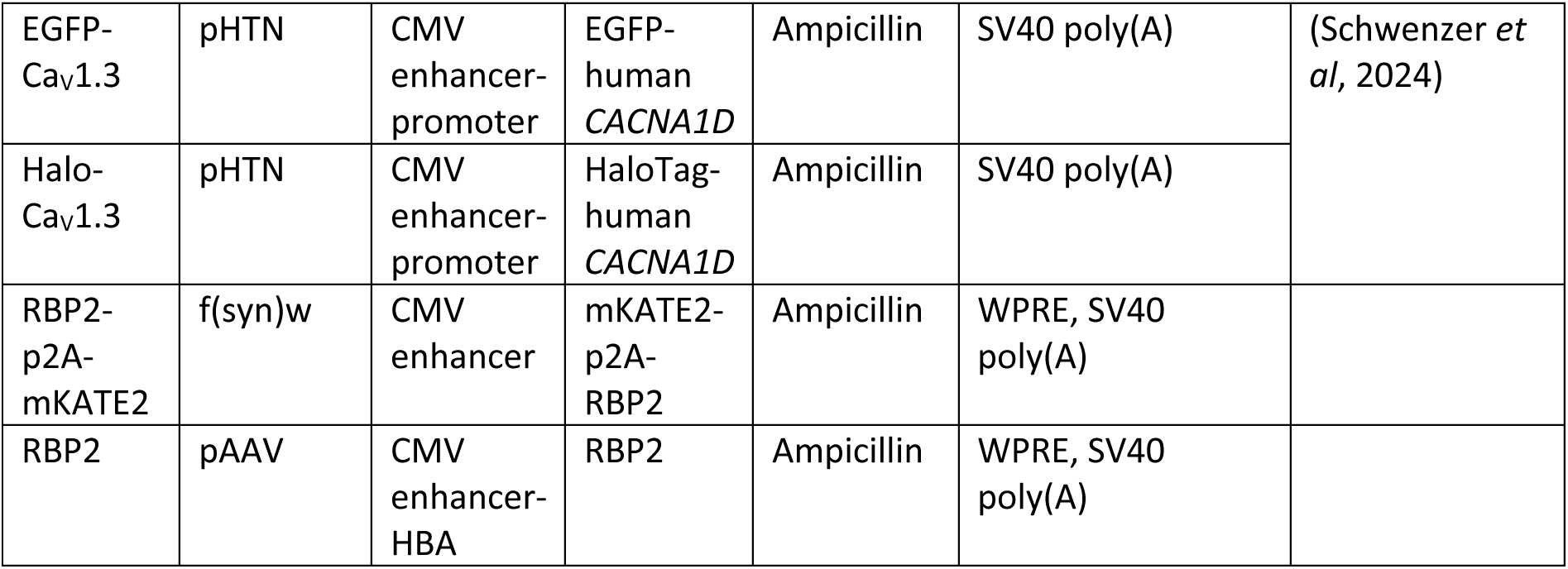
List of plasmids.

**Table 2:**
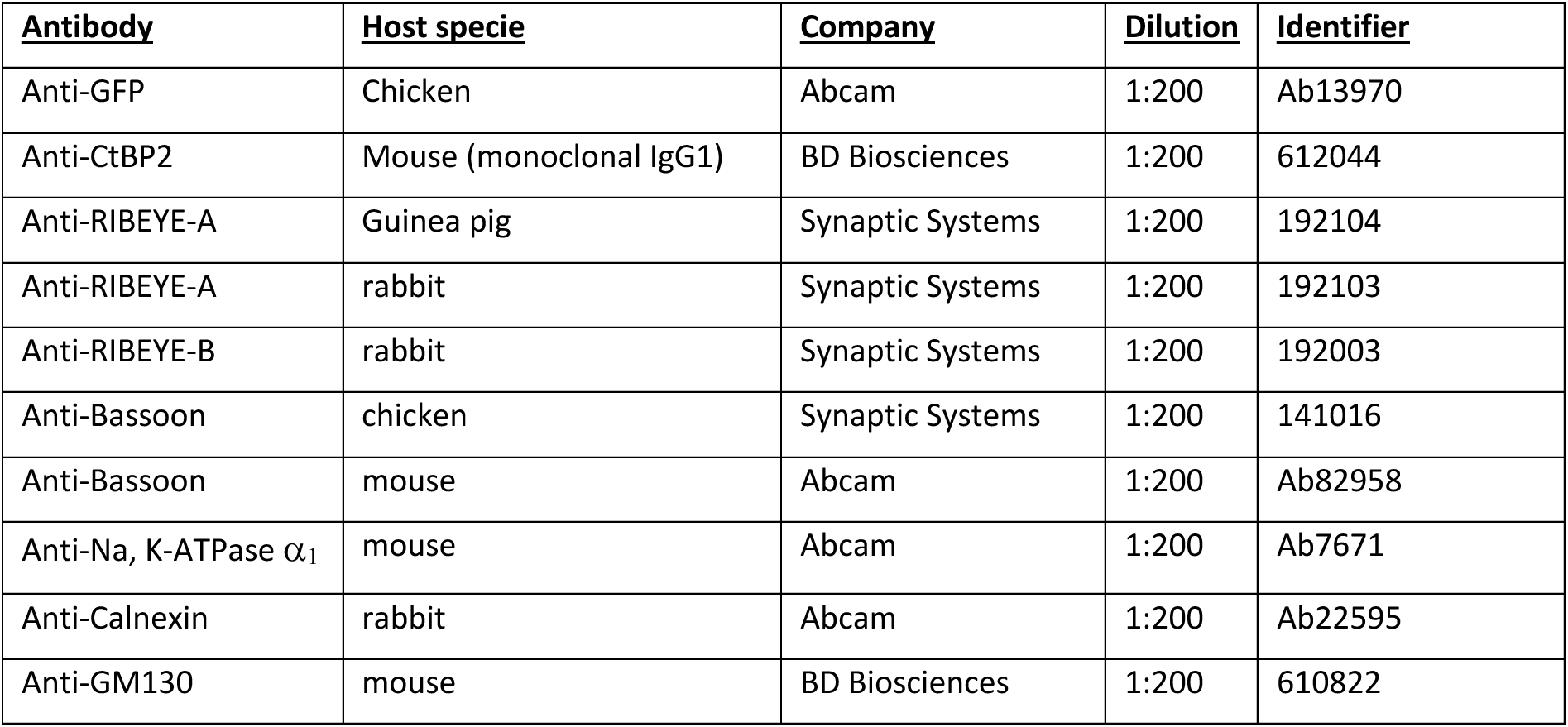
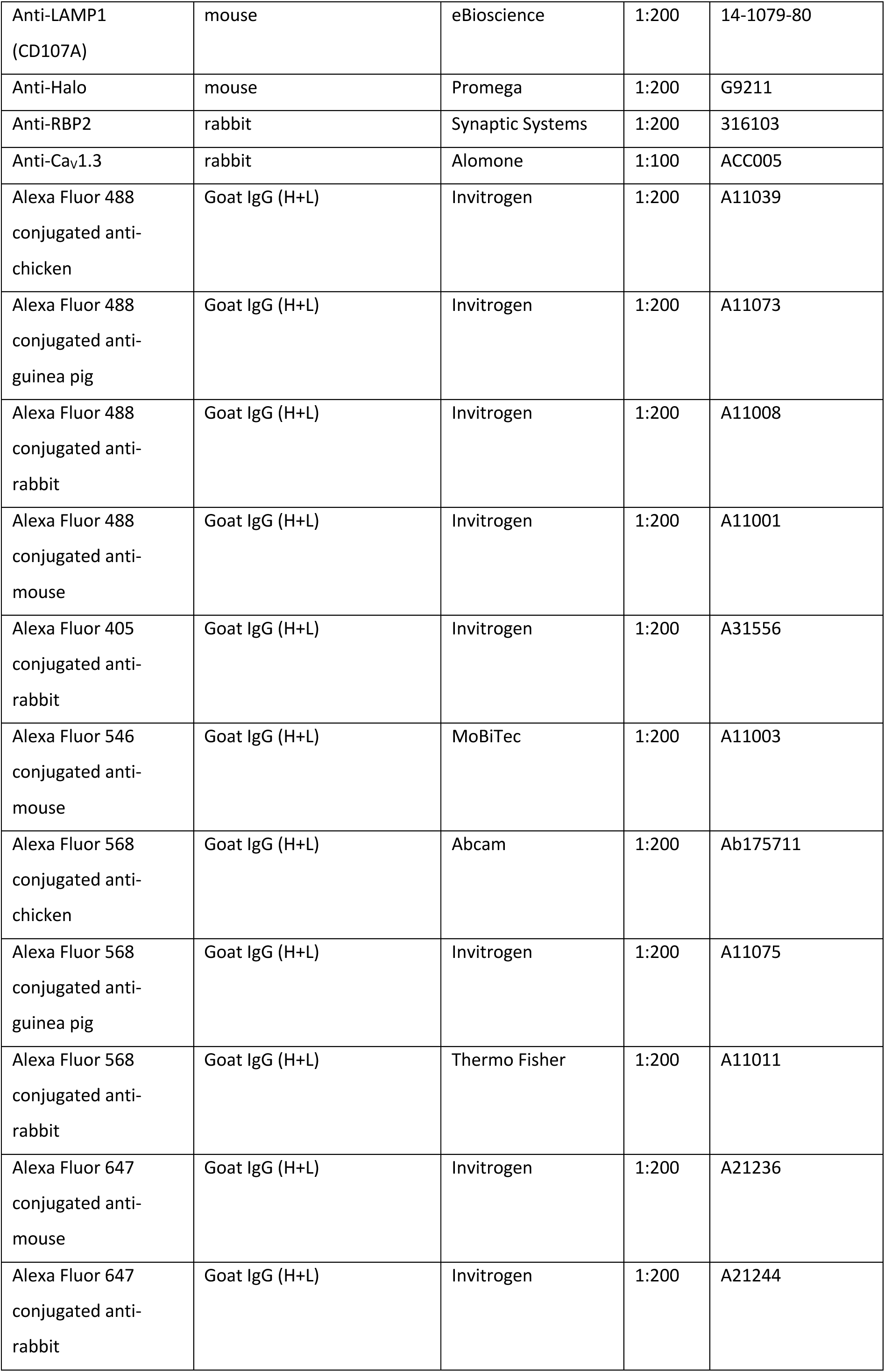
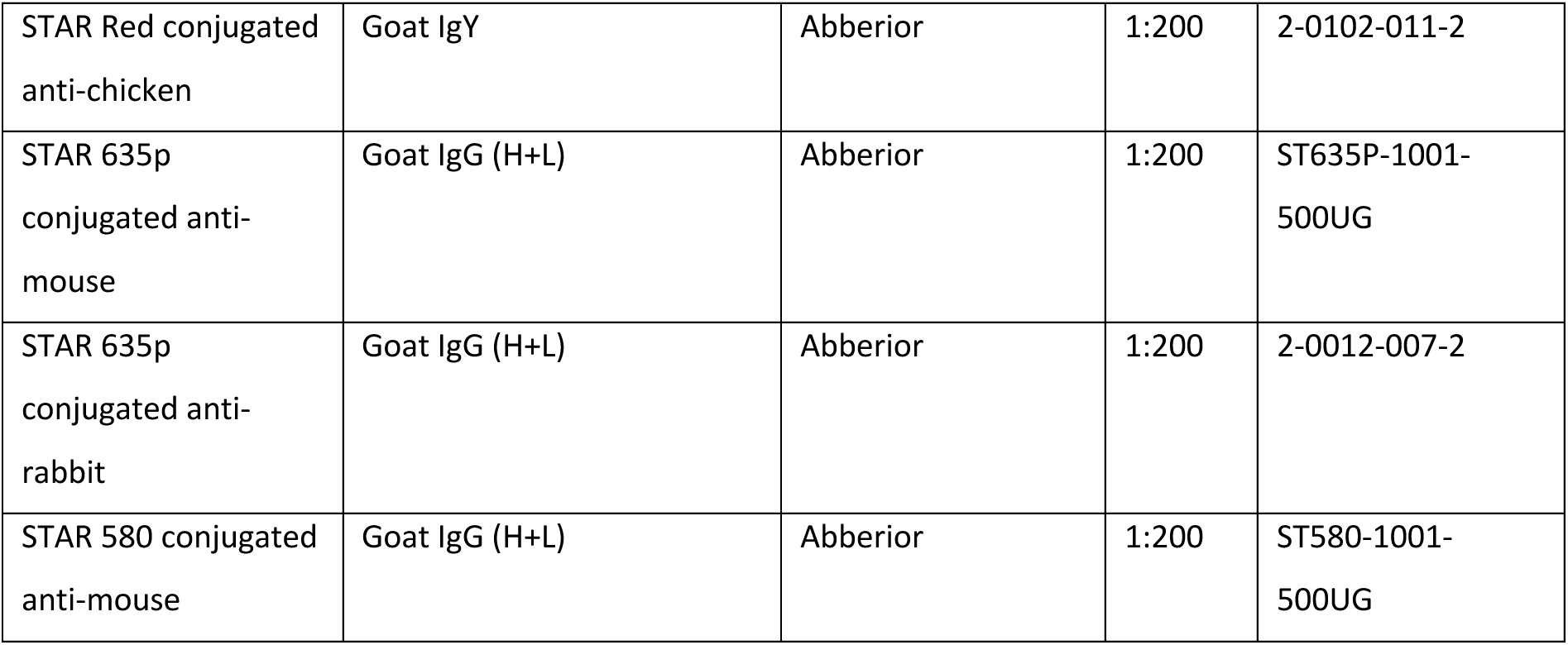
List of antibodies.

| <u>Antibody</u> | <u>Host specie</u> | <u>Company</u> | <u>Dilution</u> | <u>Identifier</u> |
| --- | --- | --- | --- | --- |
| Anti-GFP | Chicken | Abcam | 1:200 | Ab13970 |
| Anti-CtBP2 | Mouse (monoclonal IgG1) | BD Biosciences | 1:200 | 612044 |
| Anti-RIBEYE-A | Guinea pig | Synaptic Systems | 1:200 | 192104 |
| Anti-RIBEYE-A | rabbit | Synaptic Systems | 1:200 | 192103 |
| Anti-RIBEYE-B | rabbit | Synaptic Systems | 1:200 | 192003 |
| Anti-Bassoon | chicken | Synaptic Systems | 1:200 | 141016 |
| Anti-Bassoon | mouse | Abcam | 1:200 | Ab82958 |
| Anti-Na, K-ATPase $\alpha_1$ | mouse | Abcam | 1:200 | Ab7671 |
| Anti-Calnexin | rabbit | Abcam | 1:200 | Ab22595 |
| Anti-GM130 | mouse | BD Biosciences | 1:200 | 610822 |
| Anti-LAMP1<br>(CD107A) | mouse | eBioscience | 1:200 | 14-1079-80 |
| Anti-Halo | mouse | Promega | 1:200 | G9211 |
| Anti-RBP2 | rabbit | Synaptic Systems | 1:200 | 316103 |
| Anti-Cav1.3 | rabbit | Alomone | 1:100 | ACC005 |
| Alexa Fluor 488<br>conjugated anti-chicken | Goat IgG (H+L) | Invitrogen | 1:200 | A11039 |
| Alexa Fluor 488<br>conjugated anti-<br>guinea pig | Goat IgG (H+L) | Invitrogen | 1:200 | A11073 |
| Alexa Fluor 488<br>conjugated anti-rabbit | Goat IgG (H+L) | Invitrogen | 1:200 | A11008 |
| Alexa Fluor 488<br>conjugated anti-mouse | Goat IgG (H+L) | Invitrogen | 1:200 | A11001 |
| Alexa Fluor 405<br>conjugated anti-rabbit | Goat IgG (H+L) | Invitrogen | 1:200 | A31556 |
| Alexa Fluor 546<br>conjugated anti-mouse | Goat IgG (H+L) | MoBiTec | 1:200 | A11003 |
| Alexa Fluor 568<br>conjugated anti-chicken | Goat IgG (H+L) | Abcam | 1:200 | Ab175711 |
| Alexa Fluor 568<br>conjugated anti-<br>guinea pig | Goat IgG (H+L) | Invitrogen | 1:200 | A11075 |
| Alexa Fluor 568<br>conjugated anti-rabbit | Goat IgG (H+L) | Thermo Fisher | 1:200 | A11011 |
| Alexa Fluor 647<br>conjugated anti-mouse | Goat IgG (H+L) | Invitrogen | 1:200 | A21236 |
| Alexa Fluor 647<br>conjugated anti-rabbit | Goat IgG (H+L) | Invitrogen | 1:200 | A21244 |
| STAR Red conjugated anti-chicken | Goat IgY | Abberior | 1:200 | 2-0102-011-2 |
| STAR 635p conjugated anti-mouse | Goat IgG (H+L) | Abberior | 1:200 | ST635P-1001-500UG |
| STAR 635p conjugated anti-rabbit | Goat IgG (H+L) | Abberior | 1:200 | 2-0012-007-2 |
| STAR 580 conjugated anti-mouse | Goat IgG (H+L) | Abberior | 1:200 | ST580-1001-500UG |

Images from fixed and live samples were acquired in confocal/ STED mode using an Olympus IX83 inverted microscope combined with an Abberior Instruments Expert Line STED microscope (Abberior Instruments GmbH). We used lasers at 488, 561, and 633 nm for excitation and a 775 nm (1.2W) laser for stimulated emission depletion. 1.4 NA 100X or 20X oil immersion objectives were used for fixed samples, and a water immersion 60X objective was used for live samples. Confocal stacks or single sections were acquired using *Imspector* Software (pixel size = 80 X 80 nm along xy, 200 nm along z). For 2D-STED images, a pixel size of 15 X 15 nm (in xy) was used. Images in Supplementary Fig. 2A were acquired using a Leica SP8 confocal microscope (Leica Microsystems, Germany). All acquired images were visualized using NIH ImageJ software and adjusted for brightness and contrast. Samples and their corresponding controls were processed in parallel using identical staining protocols, laser excitation powers and microscope settings.

### Ultrastructural analysis using cryo-correlative electron-light microscopy

HEK293 cells were transfected with RIBEYE-GFP and untagged palm-Bassoon constructs as described above. The next day, the cells were detached using Accutase (Sigma-Aldrich, A6964) and replated on electron microscopy grids (R2/2, Au 200 mesh, 100 holey SiO_2_ film; Quantifoil) at a density of 20,000 cells/grid. After 24 hours, grids were vitrified by plunge-freezing in liquid ethane/propane using a house-made manual plunger. Cryo-focused ion beam milling and cryo-correlative light and electron microscopy were performed as previously described (Rigort *et al*, 2010; Pierson *et al*, 2024). In brief, upon transferring into an Aquilos 2 focused-ion beam/scanning electron microscope (ThermoFisher Scientific), the samples were imaged with an integrated fluorescent light microscope (ThermoFischer Scientific) at 470 nm excitation wavelength to locate cells expressing peripheral GFP, which corresponded to membrane-localised *SyRibbons*. Milling was performed at a 9° angle at these areas, using ion currents between 500 nA and 100 nA. During milling, the GFP fluorescence was monitored additionally when the lamellae reached 2 µm and 150 nm thickness to ensure that the structures of interest were captured. All fluorescent data is shown as maximum intensity projections of 4 µm z-stacks. For visualization purposes, the fluorescent image shown in Fig. 3A was corrected using the rolling ball background subtraction algorithm implemented in Fiji (Schindelin *et al*, 2012; Sternberg, 1983). After milling, lamellae were transferred into a Krios G4 cryo-transmission electron microscope (cryo-TEM; ThermoFisher Scientific). Correlation of fluorescence on 150 nm lamellae and 8700x TEM overview revealed electron-dense structures at GFP puncta. Tilt series of these densities were collected at 33k magnification using a dose-symmetric scheme (from −45° to +63°, 9° pretilt, 3° increments) and with a 20 eV energy filter. The total electron dose was 120 e^-^/Å^2^. The data was acquired using SerialEM (Mastronarde, 2005) running PACEtomo script (Eisenstein *et al*, 2023). Data pre-processing and tomogram reconstruction from tilt series were done automatically using an in-house script implementing MotionCor2, CTFFIND4 and IMOD (https://github.com/rubenlab/snartomo; Zheng *et al*, 2017; Rohou & Grigorieff, 2015; Mastronarde, 1997). To enhance contrast, a Wiener-like filter was applied (https://github.com/dtegunov/tom_deconv) on the tomograms.

### Patch clamp and calcium imaging

Ca_V_1.3 stably expressing HEK293 cells were plated on coverslips till about 40-50% confluency was achieved, following which they were induced for Ca_V_1.3α1-D expression by incubating them for 24-48 hours in induction media supplemented with tetracycline. For transfected cells, transfection was performed after 24hrs of induction in the induction media as explained above. Cells were recorded in extracellular solution containing (in mM): 150 choline-Cl, 10 HEPES, 1 MgCl_2_ and 10 CaCl_2_; pH was adjusted to 7.3 with CsOH and osmolality was 300-310 mOsm/kg. For electrophysiological recordings in Supplementary Fig. 5 and 6, pipette solution contained (in mM): 140 NMDG, 10 NaCl, 10 HEPES, 5 EGTA, 3 Mg-ATP and 1 MgCl_2_. pH was adjusted to 7.3 with methanesulfonic acid and osmolality was around 290 mOsm/kg. For Ca^2+^ imaging in Fig. 7, the pipette solution contained (in mM): 111 Cs-glutamate, 1 MgCl_2_, 1 CaCl_2_, 10 EGTA, 13 TEA-Cl, 20 HEPES, 4 Mg-ATP, 0.3 Na-GTP, 1 L-glutathione, 0.1 Calbryte590 (AAT Bioquest, Cat. No. 20706); pH was adjusted to 7.3, osmolality to 290 mOsm/kg. Resistance of patch pipettes was 3–7 MΩ. HEK293 cells were rupture patch clamped with EPC-10 amplifier (HEKA electronics, Germany) controlled by *Patchmaster* software at room temperature, as described previously (Picher *et al*, 2017b, 2017a). Cells were kept at a holding potential of −91.2 or −88 mV. All voltages were corrected for liquid junction potential offline (21.2 mV or 18 mV). Currents were leak corrected using a p/10 protocol. Recordings were discarded when leak current exceeded −50 pA, R_s_ exceeded 15 MΩ or offset potential fluctuated more than 5 mV. Current voltage (IV) relations were recorded around 1 minute after rupturing the cell, by applying increasing step-depolarisation pulses of voltage ranging from −86.2 mV to 58.8 mV, in steps of 5 mV. Depolarisation pulses of 500 ms were used for measure Ca_V_1.3 inactivation in Supplementary Fig. 6.

For Ca^2+^ imaging experiments, we used a Yokogawa CSU-X1A spinning-disk confocal scanner, mounted on an upright microscope (Zeiss Examiner) with a 63X, 1.0 NA objective (W Plan-Apochromat, Zeiss). Images were acquired using an Andor Zyla sCMOS 4.2 camera, controlled by *VisiView* 5.0 software (Visitron Systems GmbH). GFP, mKATE2 and the low-affinity Ca^2+^ indicator Calbryte590 were excited by a VS Laser Merge System (Visitron Systems) with 488 and 561 nm laser lines. Spinning disk was set to 5000 rpm. mKATE2 positive cells showing peripheral GFP expression (indicative of RIBEYE and palm-Bassoon co-expression) were identified and used for recordings. The cell was loaded for approximately a minute with Calbryte590 dye, after which we applied step depolarizations of 20 ms from −83 to +62 mV in 5-mV step increments to measure Ca^2+^ current (density)–voltage relations. Next, a central plane of the cell was selected and the GFP fluorescence (green channel) of the cell showing *SyRibbons* was imaged. This was immediately followed by imaging of the increments of Calbryte590 fluorescence (red channel) in the same plane triggered by a depolarizing pulse to +2 mV for 500 ms (frame rate = 20 Hz). For Ca^2+^ imaging experiments, we also included occasionally occurring HEK293 cells which were electrically coupled with other neighbouring HEK293 cells. For such cells, we could not fully compensate for the slow capacitive currents and we did not use them for electrophysiological analysis.

### Data analysis

#### Analysis of confocal images

All images were processed using NIH ImageJ software to make z-projections and multi-channel composites, and figures were created using Adobe Illustrator. For analysis of membrane distribution, a region of interest (ROI) of 1 µm thickness was drawn along the periphery of the cell which was identified either using Na, K ATPase α_1_ immunofluorescence or by enhancing the contrast in images of transfected cells so that the cell boundary becomes distinguishable. Mean pixel intensity was measured along this ROI and for the area enclosed within the ROI. The ratio of the mean pixel intensity (periphery: inside) per cell was reported as an average from at least three central planes; we did not use the basal and top plane. For volumetric fits of *SyRibbons,* IHC ribbons and Ca^2+^ channel clusters, we used the inbuilt surface detection algorithm from Imaris 9.6 (Oxford Instruments). The surface detail was set to 0.16 µm and largest sphere diameters for background subtraction were set as follows: for RIBEYE/CtBP2 0.3 µm, for Bassoon 0.1 µm and for Ca_V_1.3 0.08 µm. Thresholding was performed based on the quality of immunofluorescence to ensure all discernible surfaces were detected. All surfaces with less than 10 voxels were filtered out. A manual check was done to include undetected surfaces, remove surfaces not localizing within cell of interest, and split surfaces that would occasionally be clubbed together. For analysis of *SyRibbons*, only RIBEYE surfaces colocalizing with palm-Bassoon surfaces were used for quantifications (at 0.06 µm or less). Occasionally we would detect surfaces with volumes larger than 2 µm^3^ (usually cytosolic) or smaller than 0.02 µm^3^, which were not considered for analysis. For Ca_V_1.3 cluster volumetric analysis in tetra-transfected cells, clusters classified as *+SyRibbons* were at 0.045 µm or less from palm-Bassoon positive RIBEYE surfaces. All other clusters were classified as *–SyRibbons*. Only clusters localized at the periphery of the cell were used for analysis. Pixel-based intensity correlation analysis was performed using the *Coloc 2* plugin on ImageJ (http://imagej.net/Coloc_2). Analysis was performed on 1 µm thick regions of interests marking the periphery of the cell in at least three central planes (excluding the basal and top plane). Costes method was implemented with 100 repetitions to test for statistical significance of colocalisation coefficients (Costes *et al*, 2004).

#### Analysis of patch clamp and Ca^2+^ imaging data

Analysis of electrophysiology data and figure preparation was performed using Igor Pro 6 and 7 (WaveMetrics Inc.) and final figures were compiled using Adobe Illustrator. For analyzing current density-voltage relations (IVs), the evoked Ca^2+^ current was averaged from 5 to 10 ms after depolarization start. Cells expressing current amplitudes less than −49 pA and with current density greater than −60 pA/pF were not taken into account for analysis.

Ca^2+^ imaging data analysis was performed using a customized script in ImageJ. Briefly, we performed background subtraction in time series with Calbryte590 signal and then averaged 5 frames before stimulation and subtracted these from an average of three frames during stimulation to obtain ΔF images. For visualization, the images were smoothened (3 X 3 unweighted smoothening) and a composite was created by overlaying the ΔF images and the RIBEYE-GFP channel. For analyzing ΔF/F_0_, we drew circular regions of interest (ROIs) of 2µm diameter at sites with and without *SyRibbons* in the background subtracted time series. The ΔFmax/F_0_ was calculated as the of average ΔF/F_0_ in first three frames of peak Ca^2+^ signal intensity at stimulation onset in each ROI. To minimize bias while assigning regions as with or without *SyRibbons*, ROIs were assigned in the RIBEYE-GFP channel, without looking at the corresponding Calbryte590 signal.

### Statistics

Data sorting and statistics were performed using MS Excel, *Igor Pro 7* and/or *GraphPad Prism 10*. Numerical data is represented mainly as mean ± standard deviation (SD). The normality of data was assessed using Jarque-Bera test and Kolmogorov-Smirnov test, and the equality of variances was checked using *F*-test. When comparing two samples, a two-tailed unpaired Student’s *t*-test (or paired *t*-test for Fig. 7J) was performed for normally distributed samples with equal variances. If the conditions for normality and equality of variances were not met, an unpaired Mann-Whitney-Wilcoxon test was performed. When comparing multiple samples, a one-way ANOVA with post hoc Tukey’s test was used for normally distributed data with equal variances. For samples which were not normally distributed, we used Kruskal-Wallis test with *post hoc* Dunn’s multi-comparison correction. All box plots are depicted with crosses representing mean values, central bands indicating median, whiskers for 90/10 percentiles, boxes for 75/25 percentiles and individual data points overlaid. Non-significant differences have been indicated as n.s., while significant *P* values have been depicted as *\*P* for < 0.05, \**\*P* < 0.01, *\*\*\*P* < 0.001, and *\*\*\*\*P* < 0.0001.

## Acknowledgments

We would like to thank Dr. Kathrin Kusch for advice on cloning, Dr. Jakob Neef for critical feedback on the manuscript, and Nare Karagulyan for valuable discussions regarding Ca^2+^ imaging. We are grateful to Sandra Gerke, Christiane Senger-Freitag, Sina Langer, Fabienne Hahn and Ina Preuss for expert technical assistance and Patricia Räke-Kügler for the administrative support during this study. We would also like to thank Prof. Erwin Neher and Prof. Silvio Rizzoli for their feedback throughout the course of this study. Lastly, we would like to thank lab rotation students Gantavya Arora and Ugur Coskun for their initial contributions to the project.

## Author Contributions

TM and RK designed the study. RK performed the experiments and the analysis under supervision of TM. TTD performed cryo-CLEM experiments under supervision of AP and RFB. TD, NS and SEL prepared and characterized expression plasmids for membrane targeted Bassoon (TD) and EGFP/Halo-tagged Ca_V_1.3α1 (NS, SEL). RK and TM prepared the manuscript with contribution of all authors.

