## Supplementary Figures for "Establishing synthetic ribbon-type active zones in a heterologous expression system"

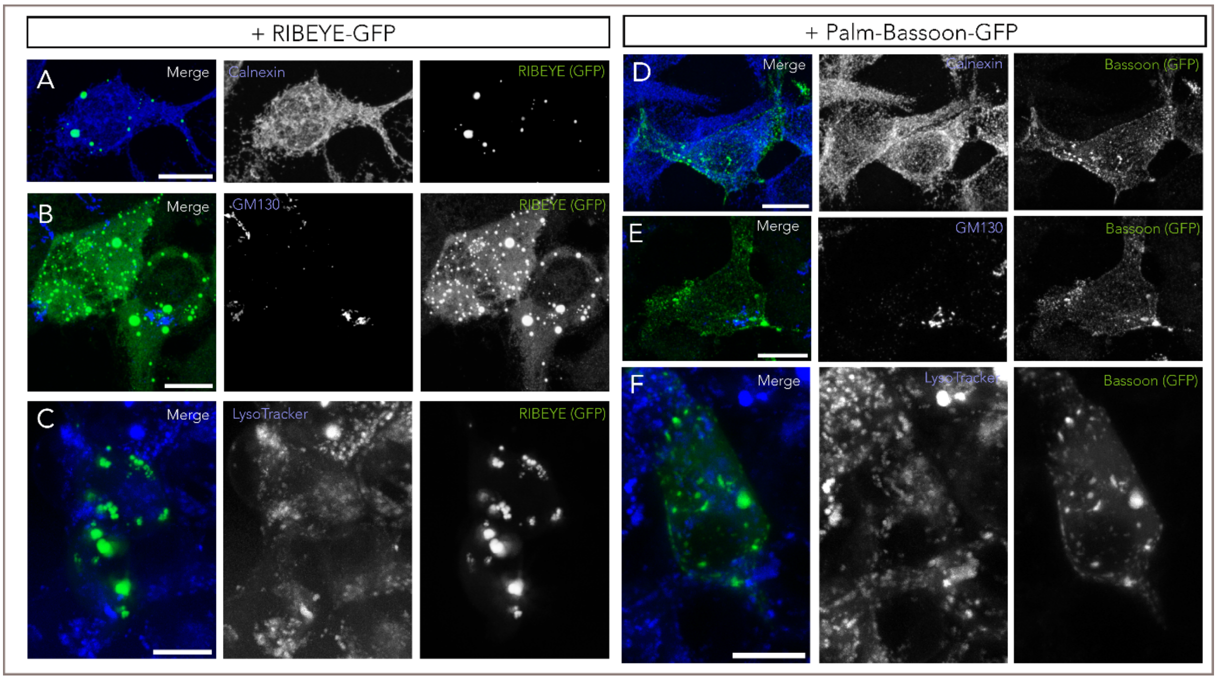


**Supplementary Figure 1 - Subcellular distribution of RIBEYE and palm-Bassoon clusters in HEK293 cells.**
Maximal projections of confocal sections of HEK293 cells transfected with RIBEYE-GFP **(A, B, C)** or palm-Bassoon-GFP **(D, E, F)**; shown in green. Representative co-staining for calnexin (endoplasmic reticulum marker), GM130 (cis-Golgi marker) and LysoTracker (live labelling of lysosomes) have been shown respectively in blue. Neither RIBEYE clusters nor palm-Bassoon appear to localize in either of the three compartments. Note that (C) and (F) represent live cells, while the remaining images are from fixed cells. Scale bar = 10 µm.

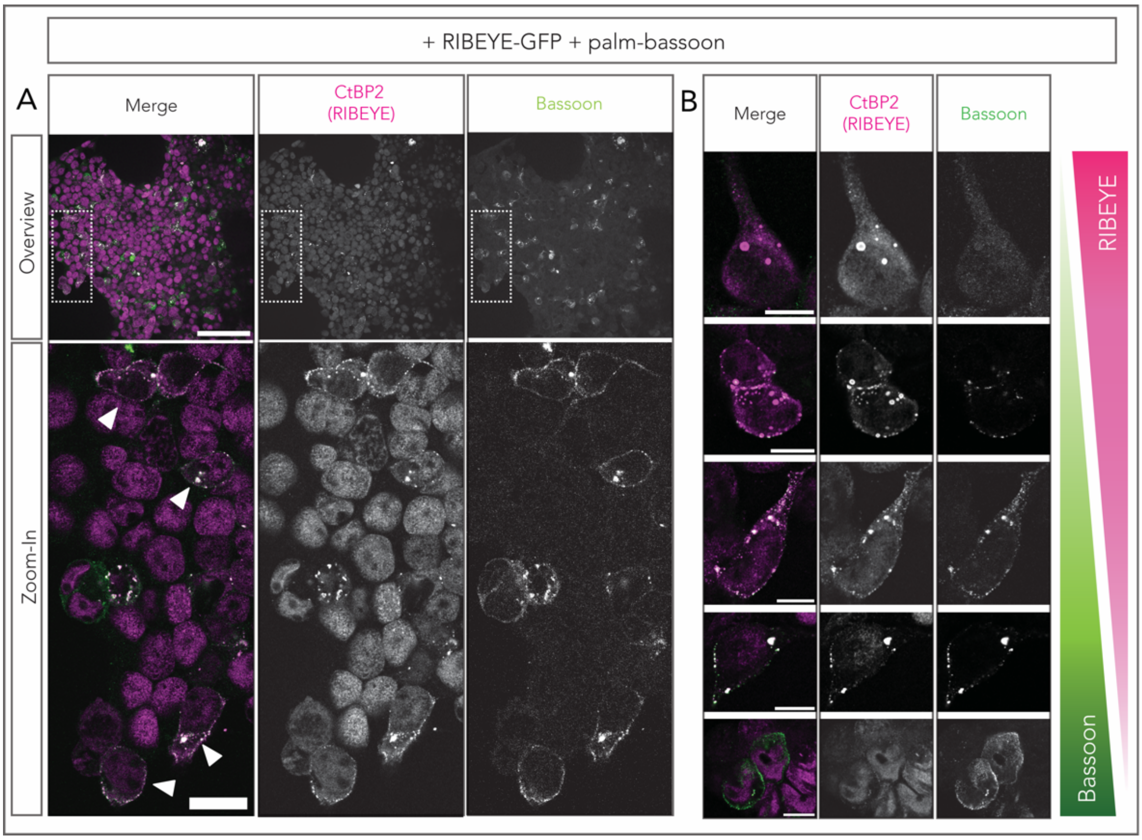


**Supplementary Figure 2 - Membrane localization of RIBEYE depends on expression levels of palm-Bassoon**

- **A** Exemplary overview image of HEK293 cells co-transfected with RIBEYE (magenta) and palm-Bassoon (green) showing how to identify cells with *SyRibbons*. On an average about 10% of cells appear co-transfected of which some show peripheral distribution of RIBEYE (examples marked by arrows). Scale bar = 100 µm for overview and 20 µm for zoom-in.
- **B** Peripheral distribution of RIBEYE appears to be dependent on expression levels of palm-Bassoon. Note the cell in the top panel with little to no palm-Bassoon expression and predominantly cytosolic RIBEYE puncta. Scale bar = 10 µm.

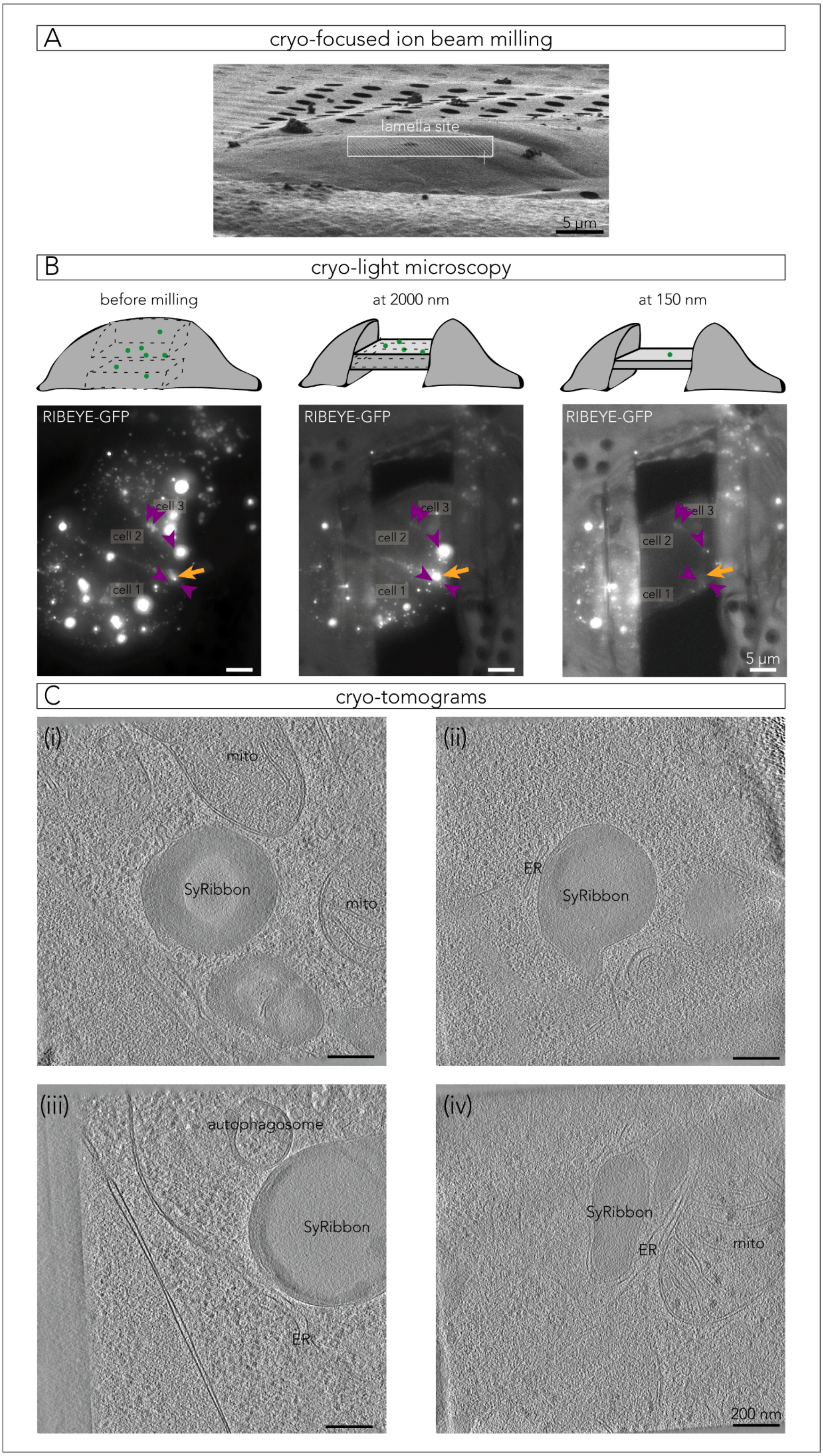

A
cryo-focused ion beam milling
lamella site
5 µm
B
cryo-light microscopy
before milling
at 2000 nm
at 150 nm
RIBEYE-GFP
cell 3
cell 2
cell 1
RIBEYE-GFP
cell 3
cell 2
cell 1
RIBEYE-GFP
cell 3
cell 2
cell 1
5 µm
C
cryo-tomograms
(i)
mito
SyRibbon
mito
(ii)
ER
SyRibbon
(iii)
autophagosome
SyRibbon
ER
(iv)
SyRibbon
ER
mito
200 nm

**Supplementary Figure 3 – Fluorescence guided milling and tomogram acquisition of *SyRibbons*.**

**A.** Vitrified group of HEK cells on an EM grid. The box indicates where a lamella was created using a cryo-focused ion beam (cryo-FIB).

**B.** Cryo-fluorescent light microscopy guided milling of HEK cells. Top row: Cartoons showing different stages of milling, where dotted lines represent regions that are ablated in the next stage. GFP fluorescence (green dots) decreases as the lamella gets thinner. Bottom row: Fluorescence acquired along the milling process. Cells expressing membrane-localised GFP are targeted for cryo-FIB milling. Here, the junction between *cell 1* and *cell 2* containing multiple GFP puncta was the region of interest. On the polished lamella (150 nm, rightmost image), a few GFP spots remained, which could often be traced back to earlier stages. In some regions, smaller puncta were occluded by bright fluorescent spots when the lamella was thick, and were only revealed at the last stage. The orange arrow indicates a peripheral signal, and purple arrowheads show cytosolic GFP.

**C.** Tomograms of *SyRibbons*. Some exhibit hollow cores (i, iii) or concentric layers (i, ii). *Mito*: mitochondrion, *ER*: endoplasmic reticulum.

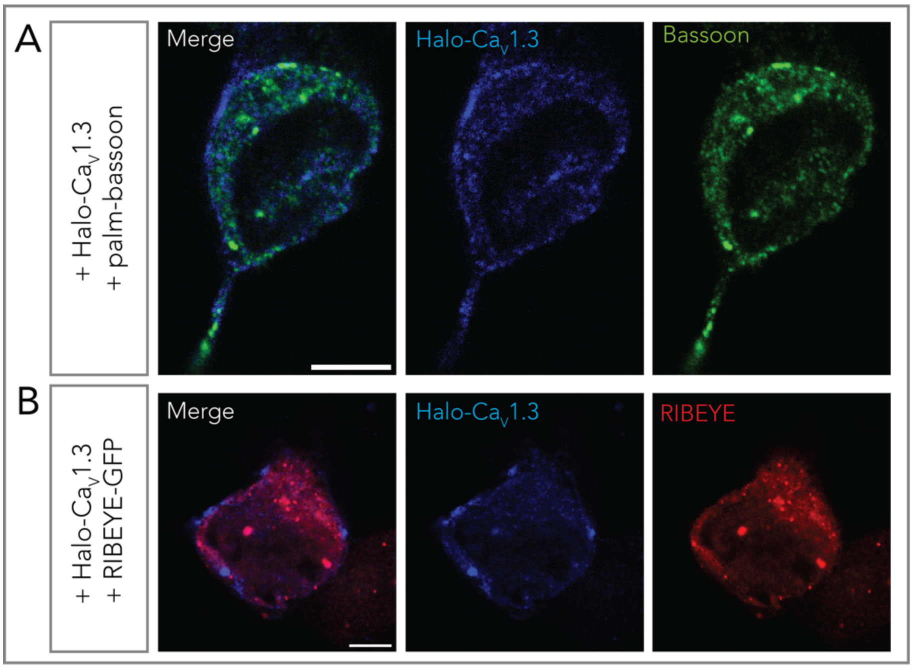


**Supplementary Figure 4 - Ca$_V$1.3 does not appear to colocalize directly with RIBEYE or with palm-Bassoon**

**A** Representative confocal section of a HEK293 cell transfected with Halo-Ca$_V$1.3 (blue) and palm-Bassoon (green). The two proteins do not appear to colocalize. Scale bar = 5 µm.

**B** Representative confocal section of a HEK293 cell transfected with Halo-Ca$_V$1.3 (blue) and RIBEYE-GFP (red). RIBEYE and Ca$_V$1.3 also do not appear to colocalize, speaking against a possible direct interaction between the two proteins. Scale bar = 5 µm.

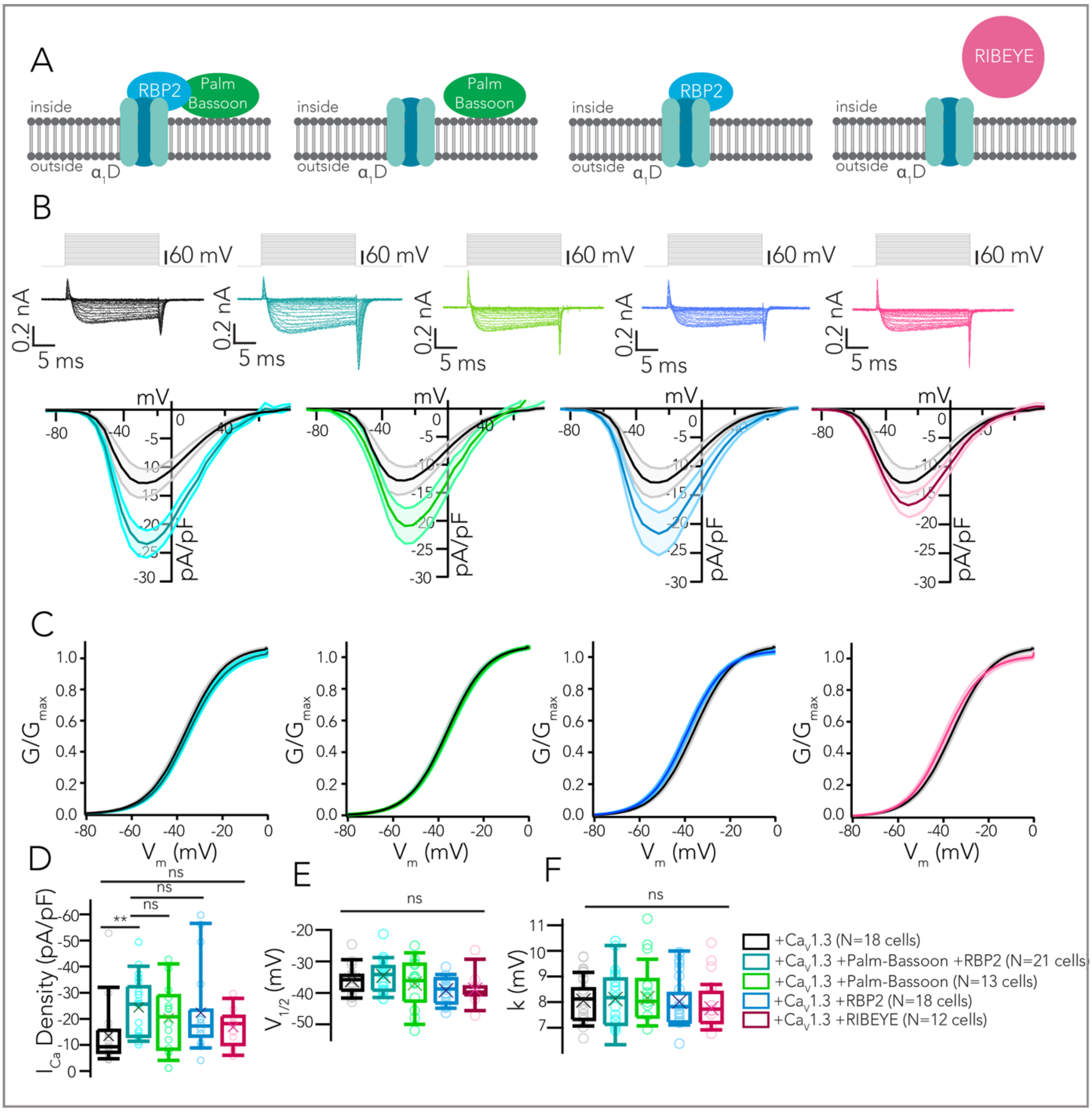

A
RBP2
Palm Bassoon
RIBEYE
inside
outside
α1D
B
60 mV
0.2 nA
5 ms
mV
pA/pF
C
G/Gmax
Vm (mV)
D
ICa Density (pA/pF)
**
ns
E
V1/2 (mV)
F
k (mV)
+Cav1.3 (N=18 cells)
+Cav1.3 +Palm-Bassoon +RBP2 (N=21 cells)
+Cav1.3 +Palm-Bassoon (N=13 cells)
+Cav1.3 +RBP2 (N=18 cells)
+Cav1.3 +RIBEYE (N=12 cells)

**Supplementary Figure 5 - Whole-cell $Ca^{2+}$ current amplitudes increase upon co-expression of RBP2 and palm-Bassoon.**

**A** Impact of co-expression of different ribbon-type AZ proteins on $Ca_V1.3\alpha_1/\beta_3/\alpha_2\delta_1$ $Ca^{2+}$ currents. Color code: non-transfected controls expressing only $Ca_V1.3$ (black), co-expression of RBP2 + palm-Bassoon (data depicted in turquoise), only palm-Bassoon (green), only RBP2 (blue) or only RIBEYE (magenta). Numbers of recorded cells/group have been indicated in the figure (bottom right).

**B** Co-expression of RBP2 and palm-Bassoon seemingly increases the whole-cell $Ca^{2+}$ current density in HEK293 cells expressing $Ca_V1.3$, $[Ca]_e$ = 10mM. Expression of palm-Bassoon or RBP2 individually also results in a mild increase, but this trend is not statistically significant as shown in (D) when compared to controls and to cells expressing both the proteins. Expression of RIBEYE by itself causes no change in $Ca^{2+}$ current density. Current density-voltage (IV) relation curves are depicted as mean $Ca^{2+}$ current density with shaded area represents $\pm$ SEM. Top panel shows representative $Ca^{2+}$ current traces.

**C** We fitted a Boltzmann function to traces in (B) to determine the fractional activation of $Ca_V1.3$ channels. The voltage-dependence of $Ca^{2+}$ current influx does not appear to be different for any of the four scenarios.

**D** Box plot depicting peak $Ca^{2+}$ current density values from (A). P values are as follows: $^{**}P_{CaV\ only/CaV+RBP2+PBsn} = 0.0041$, $P_{CaV\ only/CaV+PBsn} = 0.3737$, $P_{CaV\ only/CaV+RBP2} = 0.1643$, $P_{CaV\ only/CaV+RIBEYE} > 0.999$; Kruskal-Wallis test with *post hoc* Dunn's multiple comparison test.

**E,F** Box plots depicting voltage of half-maximal activation ($V_{half}$) and slope (k) of the Boltzmann fits in (C) show no statistically significant differences.

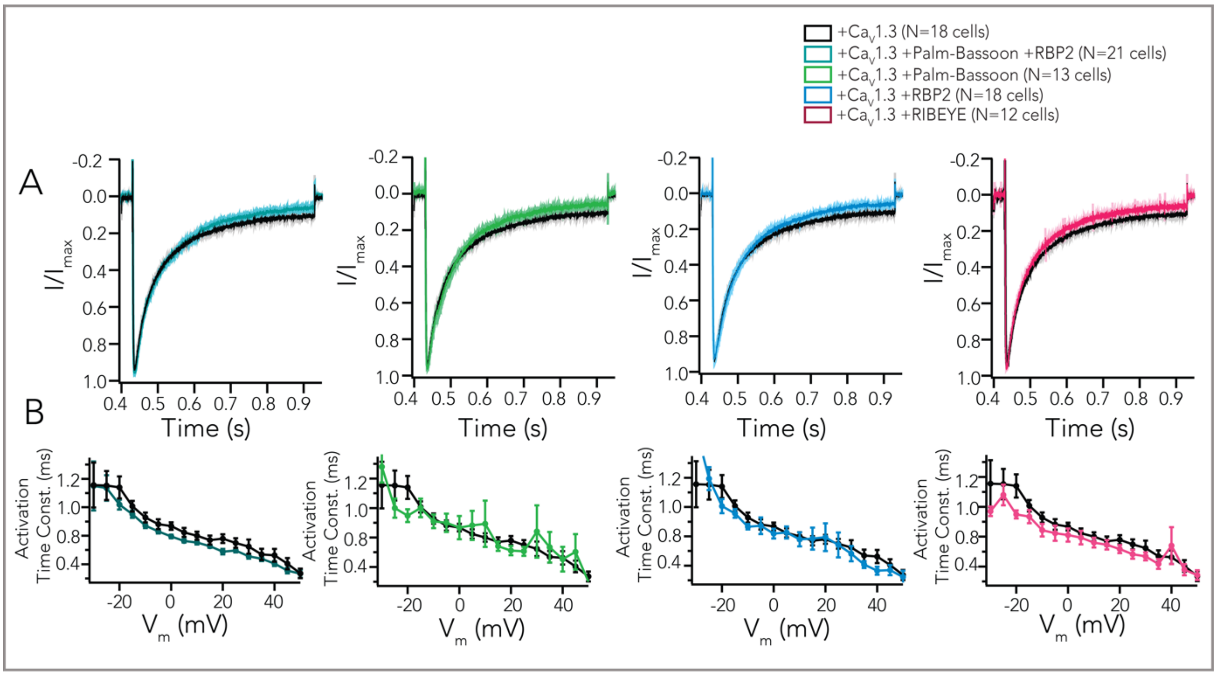


**Supplementary Figure 6 - Kinetics of Ca$_V$1.3 channels are not altered upon co-expression of various AZ proteins.**

**A** Average peak amplitude normalized $Ca^{2+}$ current traces in response to depolarizations of 500 ms from -91.2 to -21.2 mV (shaded area represents ± SEM). Expression of palm-Bassoon+RBP2 (turquoise), only palm-Bassoon (green), RBP2 (blue) or RIBEYE (magenta) do not seem to impact inactivation of Ca$_V$1.3 channels.

**B** A power exponential function was fitted to the first 5 ms of current traces (as shown in Figure 5B) to obtain the activation time constant (mean ± SEM) at different voltages, which do not appear to be notably impacted.

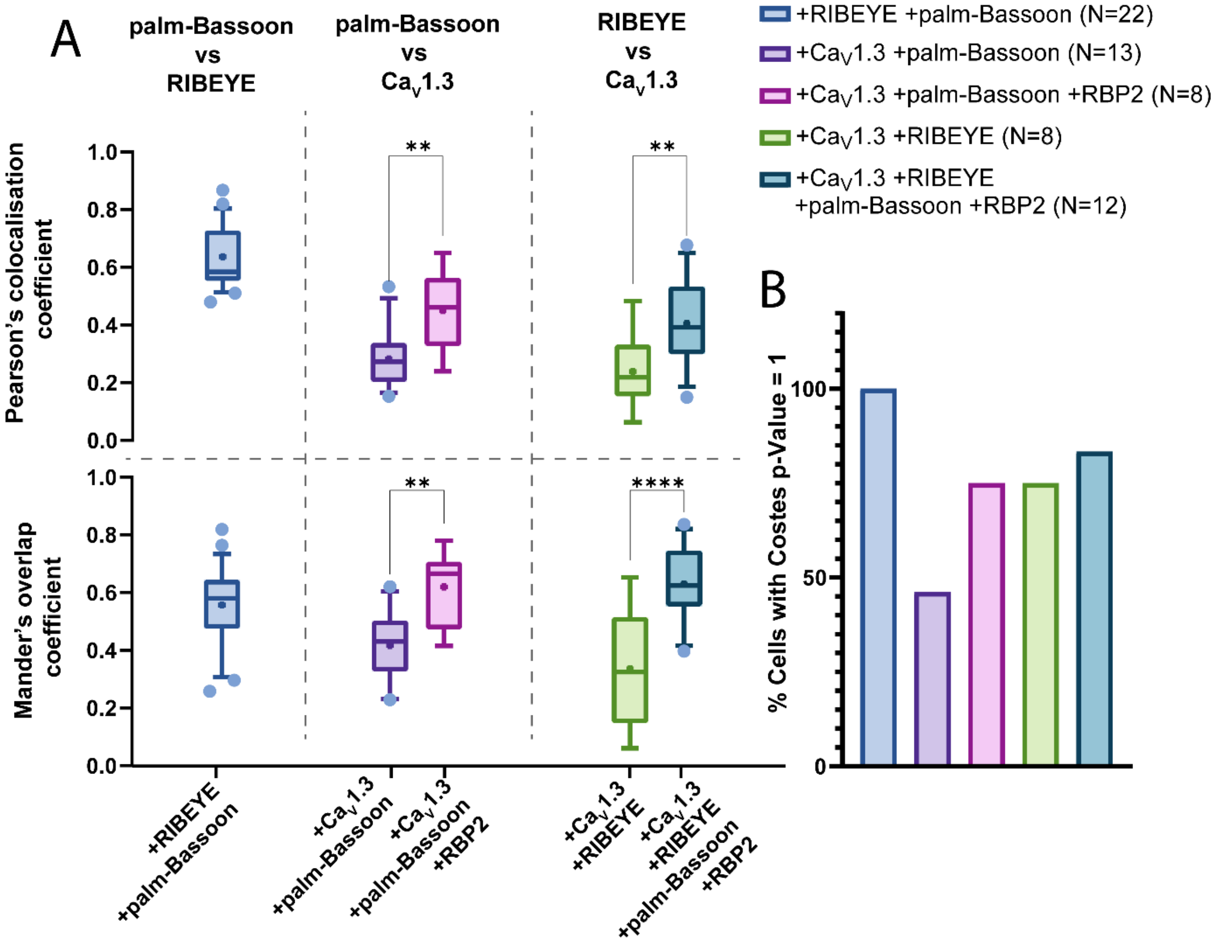


**Supplementary Figure 7 – Colocalisation quantifications**

**(A)** Box-whisker plot depicting Pearson's colocalisation coefficient and Mander's overlap coefficient between palm-Bassoon and RIBEYE, palm-Bassoon and Ca$_V$1.3 and RIBEYE and Ca$_V$1.3 in HEK293 cells transfected with the combinations of proteins described on the x-axis.

**(B)** Costes method was implemented (100 repetitions) to test for statistical significance of the determined colocalisations. The bar plot depicts percentages of cells with p = 1 indicating significant colocalisation. A p-value less than 1 indicate that the measured colocalisation is no better than random chance.
